# The Wnt pathway scaffold protein Axin promotes signaling specificity by suppressing competing kinase reactions

**DOI:** 10.1101/768242

**Authors:** Maire Gavagan, Erin Fagnan, Elizabeth B. Speltz, Jesse G. Zalatan

## Abstract

GSK3β is a multifunctional kinase that phosphorylates β-catenin in the Wnt signaling network and also acts on other protein targets in response to distinct cellular signals. To test the long-standing hypothesis that the scaffold protein Axin specifically accelerates β-catenin phosphorylation, we measured GSK3β reaction rates with multiple substrates in a minimal, biochemically-reconstituted system. We observed an unexpectedly small, ~2-fold Axin-mediated rate increase for the β-catenin reaction. The much larger effects reported previously may have arisen because Axin can rescue GSK3β from an inactive state that occurs only under highly specific conditions. Surprisingly, Axin significantly slows the reaction of GSK3β with CREB, a non-Wnt pathway substrate. When both β-catenin and CREB are present, Axin accelerates the β-catenin reaction by preventing competition with CREB. Thus, while Axin alone does not markedly accelerate the β-catenin reaction, in physiological settings where multiple GSK3β substrates are present, Axin can promote signaling specificity by suppressing interactions with competing, non-Wnt pathway targets.

## Introduction

GSK3β is a central kinase in mammalian cell signaling networks that responds to growth factors and hormones to regulate cell growth, differentiation, and metabolism (1). GSK3β receives signals from multiple upstream inputs and acts on several distinct downstream protein targets (Figure 1A). GSK3β-dependent responses to Wnt and growth factor or insulin signals appear to be insulated from each other, so that Wnt signals do not activate a growth factor/insulin response and vice versa (2–4). These observations raise the fundamental question of how GSK3β activity can be independently regulated by different signaling pathways. Analogous questions arise in many eukaryotic signaling networks, and understanding the mechanisms by which biochemical systems resolve this problem is a major outstanding challenge for the field.

**Figure 1.**
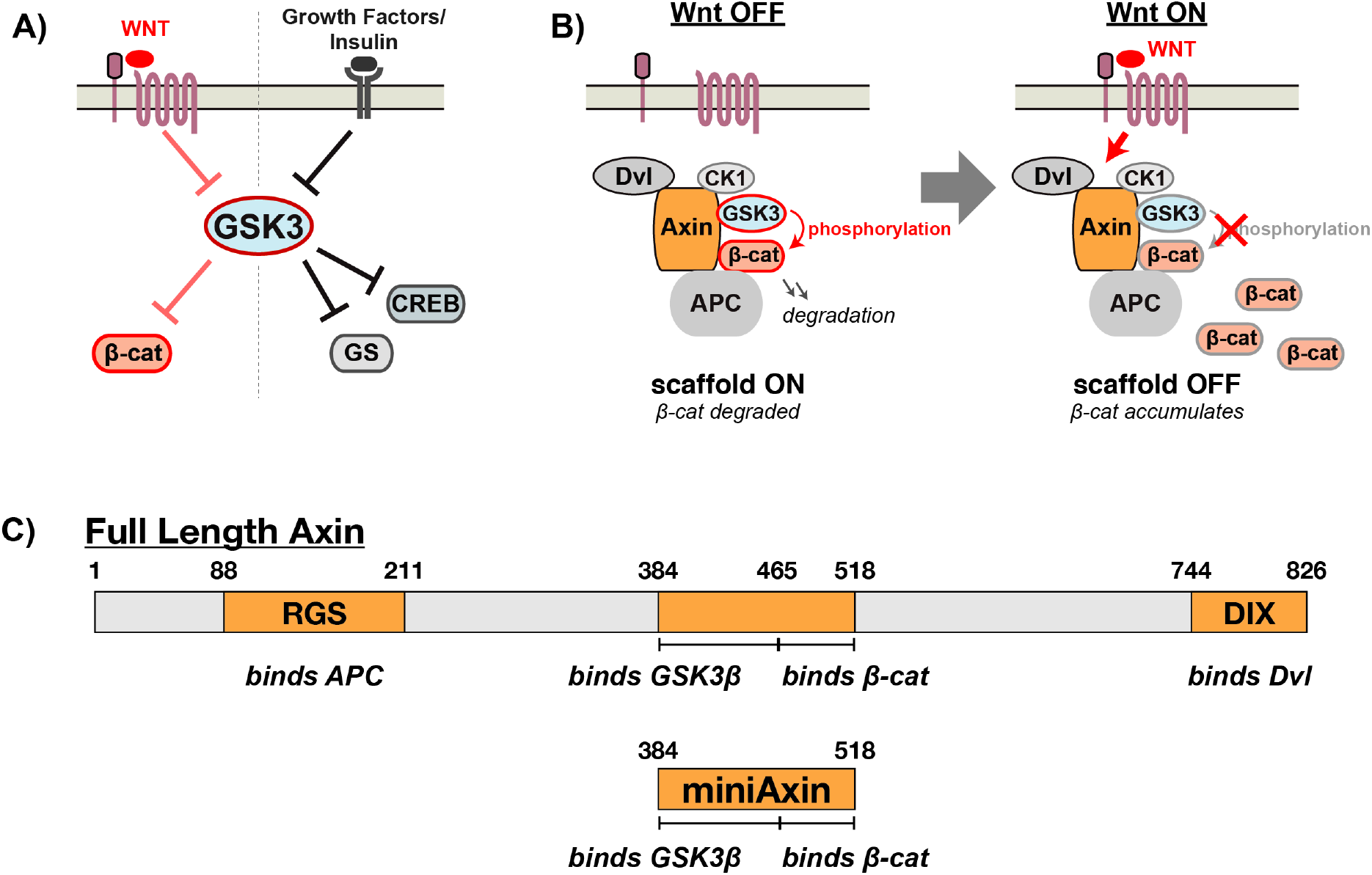
Axin assembles signaling proteins in the Wnt pathway. (A) GSK3β receives input signals from Wnt, growth factors, and hormones like insulin and acts on multiple distinct downstream targets. Wnt signals appear to be insulated from growth factor/insulin signaling. (B) GSK3β assembles into a multi-protein destruction complex with β-catenin, CK1α, Axin, Dvl, and APC. In the absence of a Wnt signal, β-catenin is phosphorylated and degraded. In the presence of a Wnt signal, β-catenin phosphorylation is blocked, which allows β-catenin to accumulate. (C) Schematics of full length human Axin (isoform 2, Uniprot O15169-2) and Axin_384-518_ (miniAxin), which contains the binding sites for both GSK3β and β-catenin (see Methods for an explanation of domain boundaries).

Scaffold proteins that physically assemble protein signaling pathways provide potential mechanisms to direct shared signaling proteins to specific downstream targets (5–7). In the Wnt signaling network, the scaffold protein Axin coordinates the assembly of a multi-protein complex including GSK3β and its substrate β-catenin. By binding to both GSK3β and β-catenin, Axin is thought to promote β-catenin phosphorylation (8–13). Consequently, regulation of Axin provides a possible mechanism to control GSK3β activity towards β-catenin without affecting GSK3β reactions towards other, non-Wnt pathway substrates (14).

We know a great deal about the general features of Wnt pathway activation, although the molecular mechanism by which Wnt signaling controls β-catenin phosphorylation and the precise role of Axin in this process remained debated (13). In the absence of a Wnt signal, β-catenin is sequentially phosphorylated by the kinases CK1α and GSK3β, which leads to proteasomal degradation of β-catenin (Figure 1B). Phosphorylation takes place in a multi-protein destruction complex that includes the scaffold protein Axin, the kinases CK1α and GSK3β, the substrate β-catenin, and the accessory proteins Dvl and APC, which may also have scaffolding functions. Wnt pathway activation recruits the destruction complex to the membrane and disrupts its activity, allowing β-catenin to accumulate and activate downstream gene expression. Wnt signaling has been proposed to block β-catenin phosphorylation at both the CK1α and GSK3β kinase reaction steps (15), possibly via a phosphorylation-dependent conformational change in Axin (16) or by inhibition of GSK3β when the destruction complex is recruited to the membrane (17).

Initial support for the model that Axin promotes β-catenin phosphorylation came from *in vitro* biochemical experiments showing that the rate of GSK3β-catalyzed β-catenin phosphorylation increases substantially in the presence of Axin (18–20). To test this hypothesis, we biochemically reconstituted GSK3β-catalyzed reactions *in vitro* and quantitatively measured reaction rates in the presence and absence of the Axin scaffold protein. By defining a minimal kinetic framework and systematically measuring rate constants for individual reaction steps, we expected to determine whether Axin promotes binding between GSK3β and β-catenin or whether Axin allosterically modulates the activity of a GSK3β•β-catenin complex. Both mechanisms have been observed with other kinase signaling scaffolds (5), and distinguishing between these possibilities is an important step towards understanding how Wnt signals might perturb Axin to regulate β-catenin phosphorylation.

Surprisingly, we found that Axin has small, ~2-fold effects on the steady state rate constants for β-catenin phosphorylation *in vitro*. We observed similar effects with CK1α-phosphoprimed β-catenin, which is the preferred substrate of GSK3β *in vivo* (21, 22), and with unprimed β-catenin, which was used in early biochemical studies (18–20). The much larger effects from Axin reported in these earlier studies appear to occur only with unprimed β-catenin under highly specific conditions, and the physiological relevance of these conditions is uncertain. We further demonstrate that Axin has an unexpected ability to suppress GSK3β activity towards other, non-Wnt pathway substrates, and that Axin can produce a >10-fold increase in the β-catenin phosphorylation rate when a competing substrate for GSK3β is present. This effect arises because Axin disrupts binding interactions between GSK3β and its substrates while at the same time tethering β-catenin to GSK3β, so that Axin rescues the binding defect for one specific substrate. In physiological conditions where there are many potential substrates for GSK3β, the ability of Axin to suppress competing reactions provides a mechanism to specifically promote the β-catenin reaction. These findings demonstrate important biochemical features of the Wnt destruction complex and reveal a new model for how scaffold proteins can promote specific signaling reactions.

## Results

### Reconstituting a minimal destruction complex

To test the model that Axin accelerates the reaction of GSK3β with β-catenin, we biochemically reconstituted a minimal reaction system for quantitative kinetic analysis. We purified recombinant human forms of GSK3β, β-catenin, and Axin as maltose binding protein (MBP) fusions. We purified active GSK3β from *E. coli* (23) and found that it was phosphorylated on the activation loop Tyr216 (Figure S1A), as had been previously reported for GSK3β purified from insect cells (20). We purified recombinant primed, phospho-Ser45-β-catenin (pS45-β-catenin) by coexpressing β-catenin with CK1α in *E. coli* (Figure S1B). For comparison, we purified unprimed β-catenin expressed in the absence of CK1α. Finally, we purified full length Axin and a minimal fragment of Axin (miniAxin, residues 384-518) that includes the domains that bind both GSK3β and β-catenin (20, 24) (Figure 1C).

To confirm that recombinant Axin binds GSK3β and β-catenin in our *in vitro* system, we performed quantitative binding assays using bio-layer interferometry. We determined that full length Axin has a *K*_D_ of 7.5 nM for GSK3β. The miniAxin fragment has a *K*_D_ of 16 nM for GSK3β (Figure S2A and Table S1). These values are similar to the previously reported *K*_D_ of 65 nM for the interaction between human GSK3β and rat Axin (18). Because miniAxin behaved similarly to full length Axin, and because this construct was easier to express and purify in larger quantity than the full-length protein, we used miniAxin for most subsequent binding and kinetic assays. For several key experiments, we verified that full length Axin gave similar behavior.

We proceeded to measure the affinity of miniAxin for pS45-β-catenin. The observed *K*_D_ of 4.0 µM (Figure S2B and Table S1) is similar to the reported *K*_D_ of 1.6 µM for unprimed mouse β-catenin binding to a short human Axin fragment (residues 436-498) (25). We also attempted to measure an affinity with unprimed β-catenin. We could detect miniAxin binding to unprimed β-catenin in a similar concentration range, but we were unable to accurately measure a *K*_D_ value due to surface aggregation artifacts. Taken together, we confirmed that Axin binds to both GSK3β and β-catenin, and that Axin binds GSK3β substantially more tightly than β-catenin.

To determine how Axin affects the β-catenin phosphorylation reaction, we directly compared the steady state rate constants *k*_cat_/*K*_M_ and *k*_cat_ in the presence and absence of Axin (Figure 2A). Because Axin binds GSK3β much more tightly than β-catenin, we can define a minimal, simplified kinetic scheme that allows straightforward comparisons. We worked at a fixed Axin concentration above the *K*_D_ for GSK3β but below the *K*_D_ for β-catenin, which ensures that all GSK3β is bound to Axin. Although there is excess free Axin in the system, the concentration of Axin is below the *K*_D_ for β-catenin and there should be very little β-catenin bound to free Axin. By varying the β-catenin concentration, we can obtain the steady state rate constants for the Axin•GSK3β complex, which can be compared to the values obtained with GSK3β in the absence of Axin. We expect that if Axin promotes binding between GSK3β and β-catenin, the observed *K*_M_ should shift to a lower concentration and *k*_cat_ should remain unchanged. Alternatively, if Axin allosterically activates the GSK3β•β-catenin complex, then *k*_cat_ should increase without necessarily affecting *K*_M_.

**Figure 2.**
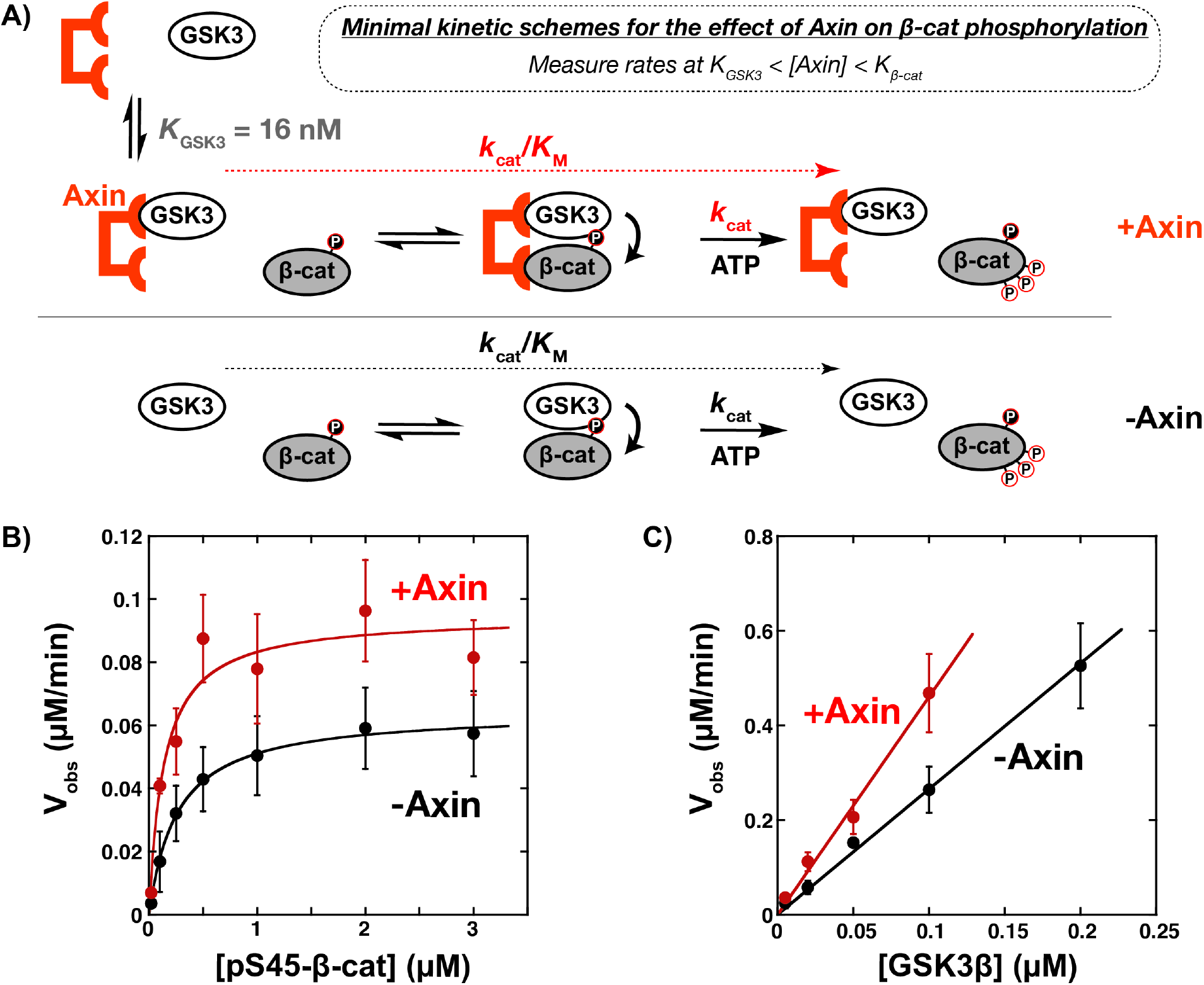
Axin has a modest effect on phosphorylation of phosphoprimed β-catenin. (A) Minimal kinetic scheme for the reaction of GSK3β with pS45-β-catenin in the presence and absence of Axin. GSK3β phosphorylates pS45-β-catenin at three sites: S33, S37, and T41. When the Axin concentration is larger than the affinity for GSK3β (*K*_GSK3_) but smaller than the affinity for pS45-β-catenin (*K*_β-cat_), all GSK3β is bound to Axin (when [Axin] > [GSK3β]). By varying the concentration of pS45-β-catenin, we can obtain the Michaelis-Menten kinetic parameters *k*_cat_, *K*_M_, and *k*_cat_/*K*_M_ for the reaction with the Axin•GSK3β complex. These parameters can be compared directly to the corresponding parameters for the reaction in the absence of Axin. (B) Michaelis-Menten plot of *V*_obs_ vs. [pS45-β-catenin] at 20 nM GSK3β in the presence and absence of 500 nM miniAxin. See Table 1 for values of fitted kinetic parameters. (C) Plot of *V*_obs_ vs. [GSK3β] at 1 µM pS45-β-catenin in the presence and absence of 500 nM miniAxin. *V*_obs_ increases linearly with enzyme concentration, as expected. An accurate initial rate for 0.2 µM GSK3β with Axin could not be measured because the reaction exceeded 40% conversion before two timepoints could be collected. Error bars in (B) & (C) are mean ± SD for at least 3 measurements.

To measure β-catenin phosphorylation rates, we used quantitative Western blotting. GSK3β sequentially phosphorylates pS45-β-catenin at three sites: T41, S37, and S33 (22), and product formation can be monitored using an antibody specific for pS33/pS37/pT41-β-catenin (see Methods). In the presence of miniAxin, we observed a 1.5-fold increase in *k*_cat_, a 2-fold decrease in *K*_M_, and a 3-fold increase in *k*_cat_/*K*_M_ (Figure 2B & C, Table 1). These effects are far smaller than expected based on prior reports, one of which suggested that Axin accelerates the reaction by >10^4^-fold (18–20).

**Table 1.**
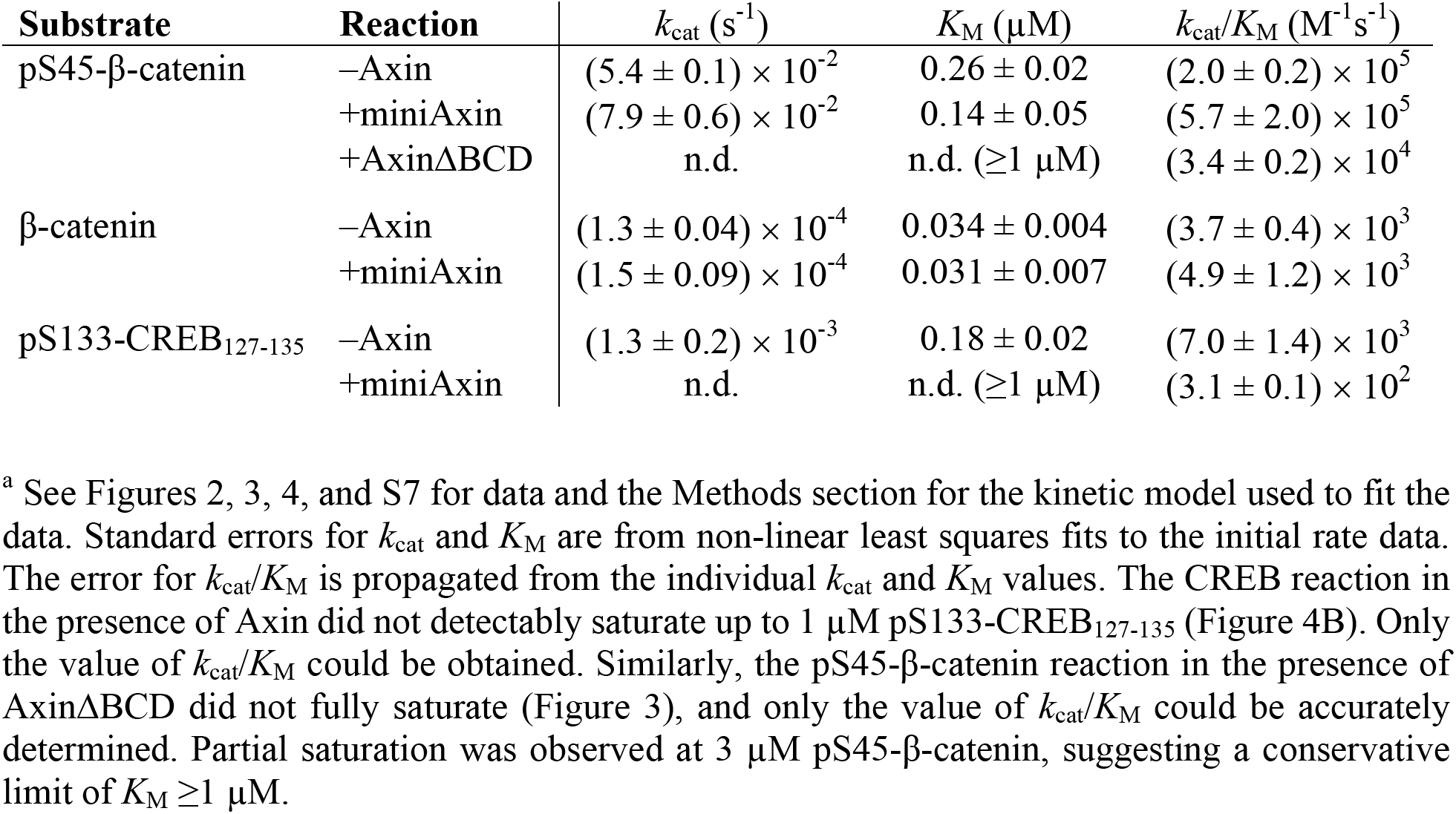
Kinetic parameters for GSK3β reactions.^a^

| Substrate | Reaction | $k_{\text{cat}}$ (s<sup>-1</sup>) | $K_M$ (μM) | $k_{\text{cat}}/K_M$ (M<sup>-1</sup>s<sup>-1</sup>) |
| --- | --- | --- | --- | --- |
| pS45-β-catenin | -Axin | $(5.4 \pm 0.1) \times 10^{-2}$ | $0.26 \pm 0.02$ | $(2.0 \pm 0.2) \times 10^5$ |
| | +miniAxin | $(7.9 \pm 0.6) \times 10^{-2}$ | $0.14 \pm 0.05$ | $(5.7 \pm 2.0) \times 10^5$ |
| | +AxinΔBCD | n.d. | n.d. ($\geq 1 \mu$M) | $(3.4 \pm 0.2) \times 10^4$ |
| β-catenin | -Axin | $(1.3 \pm 0.04) \times 10^{-4}$ | $0.034 \pm 0.004$ | $(3.7 \pm 0.4) \times 10^3$ |
| | +miniAxin | $(1.5 \pm 0.09) \times 10^{-4}$ | $0.031 \pm 0.007$ | $(4.9 \pm 1.2) \times 10^3$ |
| pS133-CREB<sub>127-135</sub> | -Axin | $(1.3 \pm 0.2) \times 10^{-3}$ | $0.18 \pm 0.02$ | $(7.0 \pm 1.4) \times 10^3$ |
| | +miniAxin | n.d. | n.d. ($\geq 1 \mu$M) | $(3.1 \pm 0.1) \times 10^2$ |

<sup>a</sup> See Figures 2, 3, 4, and S7 for data and the Methods section for the kinetic model used to fit the data. Standard errors for $k_{\text{cat}}$ and $K_M$ are from non-linear least squares fits to the initial rate data. The error for $k_{\text{cat}}/K_M$ is propagated from the individual $k_{\text{cat}}$ and $K_M$ values. The CREB reaction in the presence of Axin did not detectably saturate up to 1 μM pS133-CREB<sub>127-135</sub> (Figure 4B). Only the value of $k_{\text{cat}}/K_M$ could be obtained. Similarly, the pS45-β-catenin reaction in the presence of Axin $\Delta$ BCD did not fully saturate (Figure 3), and only the value of $k_{\text{cat}}/K_M$ could be accurately determined. Partial saturation was observed at 3 μM pS45-β-catenin, suggesting a conservative limit of $K_M \geq 1 \mu$ M.

We considered several possible explanations for the discrepancy between our results and prior work. First, scaffold-dependent reactions can be slow if too little scaffold is present, but inhibited at excess scaffold concentrations if the kinase and substrate are not bound to the same scaffold (26). While we chose the Axin concentration carefully to avoid this issue, it is possible that our assumptions were incorrect. We therefore varied the concentration of miniAxin and measured reaction rates at both saturating and subsaturating pS45-β-catenin concentrations, but found no miniAxin concentration that produced a larger rate enhancement (Figure S4). Second, it is possible that the miniAxin fragment lacks some important functional domain. When we measured reaction rates in the presence of full length Axin, however, there was no significant difference compared to miniAxin (Figure S4). Thus, Axin has only a modest effect on the GSK3β reaction with pS45-β-catenin, and this effect arises from small changes in both *k*_cat_ and *K*_M_. The decrease in *K*_M_ suggests that Axin promotes binding between GSK3β and pS45-β-catenin by a factor of ~2-fold. The effect is relatively small, possibly because Axin binding to β-catenin is weak (*K*_D_ ~4 µM) compared to the *K*_M_ for the reaction of GSK3β with pS45-β-catenin (*K*_M_ = 0.26 µM). In order to obtain large tethering effects from a scaffold, it may be necessary for binding affinities to the scaffold to be at least comparable to the un-scaffolded interaction between kinase and substrate, or for ternary complex formation to be highly cooperative.

### Removing the β-catenin binding site on Axin disrupts the activity of the Axin•GSK3β complex

If the interaction of Axin with β-catenin is weak enough that both the Axin•GSK3β complex and free GSK3β have similar *K*_M_ values for β-catenin, then the β-catenin binding site on Axin should be dispensable. To test this prediction, we removed the β-catenin binding domain (BCD) from Axin (Figure 3A). Because GSK3β binds to both Axin and Axin∆BCD with similar affinity (Table S1), we assessed the effect of Axin∆BCD on GSK3β activity using the same minimal kinetic framework as for wild type Axin (Figure 2A). In the presence of Axin∆BCD, the observed rates for phosphorylation of pS45-β-catenin were substantially slower than the corresponding rates for free GSK3β (Figure 3B), indicating that the activity of the Axin∆BCD•GSK3β complex is somehow suppressed compared to free GSK3β. The reaction does not fully saturate at high pS45-β-catenin concentrations, so we can estimate a conservative limit that *K*_M_ ≥ ~1 µM, which is significantly larger than the *K*_M_ of 0.26 µM in the absence of Axin (Table 1). A simple model to explain this observation is that Axin interferes with the ability of GSK3β to bind substrates. Based on the crystal structure of Axin bound to GSK3β, Axin does not physically occlude the substrate binding site of GSK3β (20, 27). Further, there are no obvious structural changes at this site between GSK3β bound to Axin and free GSK3β. Nevertheless, there are well-established examples of remote allosteric effects that are not detectable in static crystal structures (28). Although we lack a structural model, the kinetic data strongly suggest that Axin binding to GSK3β allosterically disrupts substrate binding. This detrimental effect is rescued by Axin binding to β-catenin, which results in the net 2-fold decrease in *K*_M_ observed for Axin•GSK3β versus GSK3β alone. We initially concluded that the 2-fold *K*_M_ decrease implied that Axin makes only a modest contribution to β-catenin binding to the Axin•GSK3β complex. However, if we compare Axin•GSK3β to Axin∆BCD•GSK3β, there is a ≥7-fold decrease in *K*_M_ (Table 1), suggesting that the β-catenin binding domain on Axin does play an important role in promoting β-catenin binding to GSK3β.

**Figure 3.**
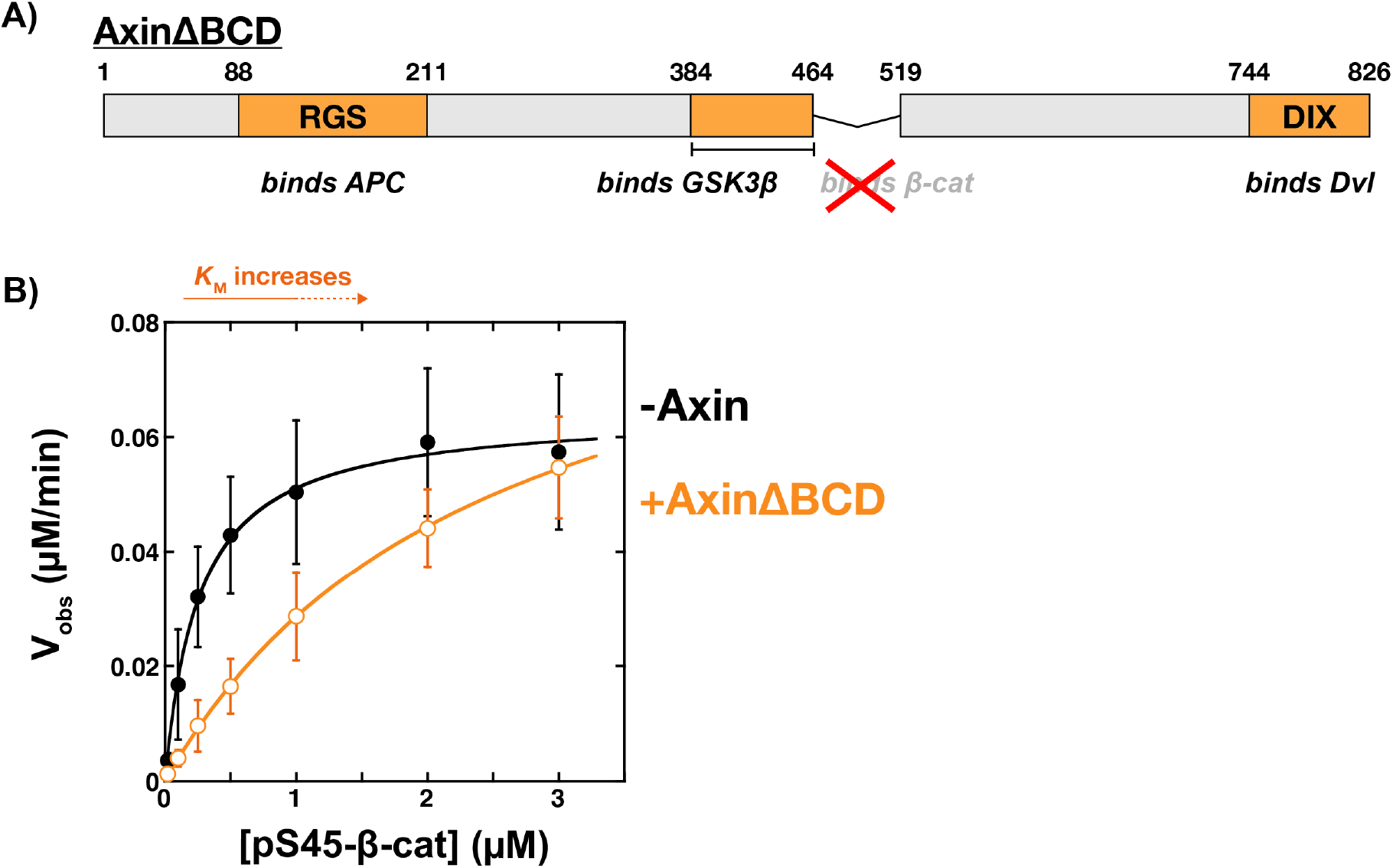
Removing the β-catenin binding site on Axin disrupts the activity of the Axin•GSKβ complex. (A) Schematic of Axin∆BCD (see Methods for explanation of domain boundaries). (B) Michaelis-Menten plot of *V*_obs_ vs. [pS45-β-catenin] at 20 nM GSK3β in the presence and absence of 500 nM Axin∆BCD. Data for the reaction in the absence of Axin is reproduced from Figure 2B. See Table 1 for values of fitted kinetic parameters. The Axin∆BCD reaction does not fully saturate at high [pS45-β-catenin], which means that only the value of *k*_cat_/*K*_M_ can be accurately determined. The reaction may be starting to saturate, suggesting a conservative limit of *K*_M_ ≥ ~1 µM.

### Axin slows the GSK3β reaction with CREB, a non-Wnt pathway substrate

If Axin has two competing functions, disrupting GSK3β substrate binding and promoting binding to β-catenin, we predicted that Axin should have a detrimental effect on the activity of GSK3β with alternative, non-Wnt pathway substrates that do not bind Axin. To determine how Axin affects interactions with these substrates, we measured reaction rates with CREB, a transcription factor that is phosphorylated by GSK3β and integrates signals from a number of pathways (29, 30). In cells, PI3K/Akt signaling represses GSK3β activity, which affects CREB-dependent transcription but does not activate Wnt outputs (4, 31).

We expressed a short CREB peptide (CREB_127-135_) as an MBP fusion; this peptide is phosphorylated by PKA at Ser133, which serves as a priming site for GSK3β to phosphorylate Ser129 (29) (Figure 4A). We phosphorylated CREB_127-135_ to completion *in vitro* with PKA to produce pS133-CREB_127-135_ (Figure S5) and measured reaction rates for GSK3β–catalyzed phosphorylation of pS133-CREB_127-135_. As predicted, the presence of Axin resulted in a substantial decrease in CREB phosphorylation rates (Figure 4B & C), with the value of *k*_cat_/*K*_M_ decreasing ~20-fold compared to the reaction without Axin (Table 1). The reaction did not detectably saturate in the presence of Axin, suggesting that the *K*_M_ has shifted to a much larger value and that at least some of the decrease in *k*_cat_/*K*_M_ arises from a disruption of binding interactions between GSK3β and pS133-CREB_127-135_. Thus, when Axin does not interact with the GSK3β substrate, Axin reduces GSK3β activity by interfering with substrate binding. This effect occurs in the CREB reaction because Axin does not have a binding site for CREB, and in the β-catenin reaction if Axin is mutated to Axin∆BCD, which lacks the β-catenin binding site.

**Figure 4.**
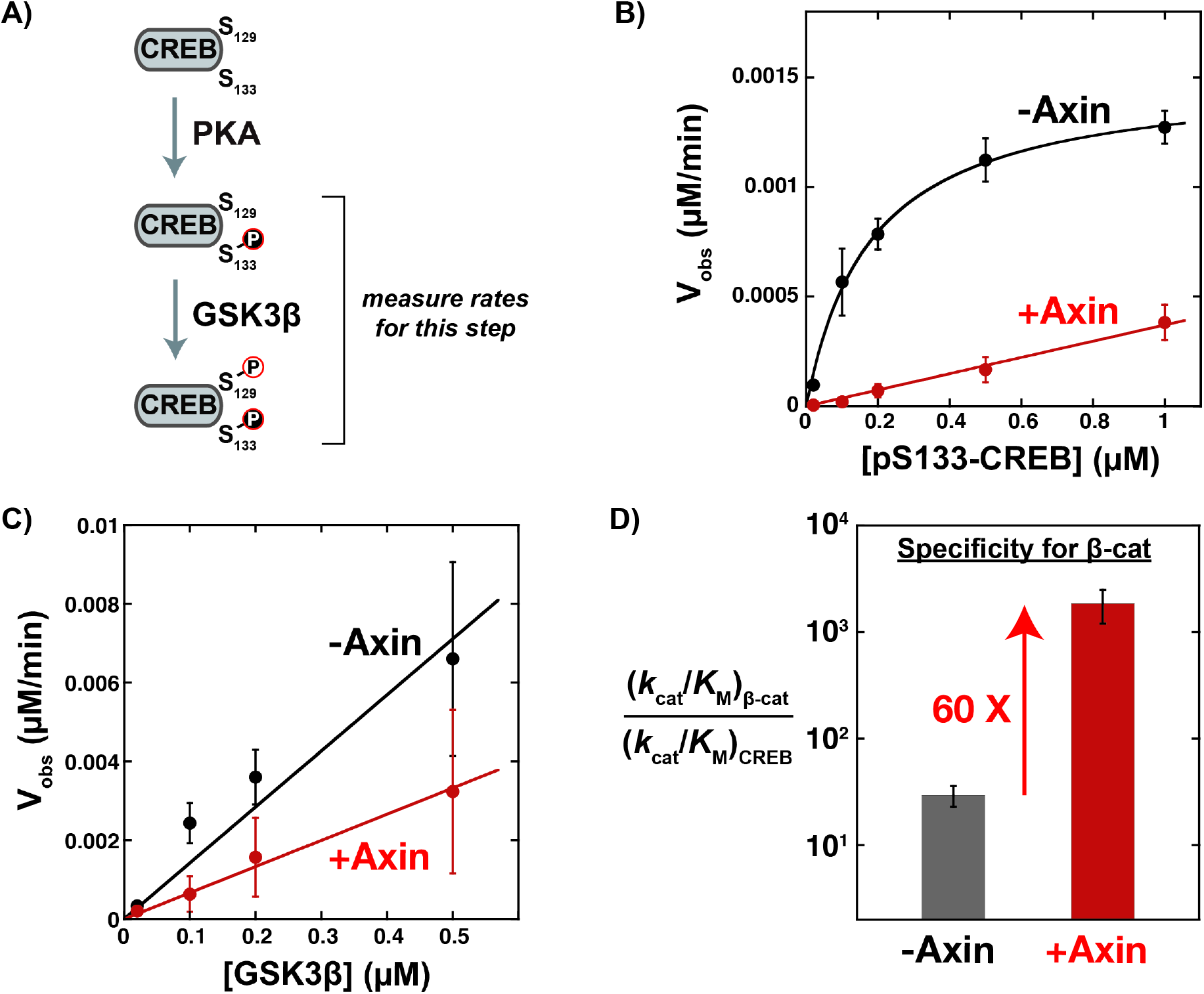
Axin significantly decreases the rate of CREB phosphorylation. (A) Minimal scheme for phosphorylation of CREB. PKA phosphorylates CREB at S133. pS133 serves as a priming site for GSK3β, which phosphorylates pS133-CREB at S129. (B) Michaelis-Menten plot of *V*_obs_ vs. [pS133-CREB_127-135_] at 20 nM GSK3β in the presence and absence of 500 nM miniAxin. See Table 1 for values of fitted kinetic parameters. There is no detectable saturation of the reaction in the presence of miniAxin up to 1 µM pS133-CREB_127-135_, which means that only the value of *k*_cat_/*K*_M_ can be determined from the slope of a linear fit to the data. We were unable to detect GSK3β-catalyzed phosphorylation of unprimed CREB (*V*_obs_ ≤ 3 × 10^−5^ µM/min at 0.5 µM CREB). (C) Plot of *V*_obs_ vs. [GSK3β] at 500 nM pS133-CREB_127-135_ in the presence and absence of 500 nM miniAxin. Error bars in (B)-(C) are mean ± SD for at least 3 measurements. (D) The specificity of GSK3β for β-catenin relative to CREB, defined as the ratio of the corresponding *k*_cat_/*K*_M_ values, increases by 60-fold in the presence of Axin. Values are from Table 1.

### Axin accelerates the β-catenin reaction when competing substrates are present

The specificity of GSK3β towards competing substrates is determined by the relative substrate concentrations and the ratio of the *k*_cat_/*K*_M_ values (Figure S6). The 20-fold drop in *k*_cat_/*K*_M_ towards pS133-CREB in the presence of Axin (Figure 4), together with the 3-fold increase in *k*_cat_/*K*_M_ towards pS45-β-catenin (Figure 2), means that the Axin•GSK3β complex is ~60-fold more specific towards β-catenin than free GSK3β (Figure 4D). When both β-catenin and CREB are present at low, subsaturating concentrations, this specificity increase from Axin will manifest largely as a decrease in the rate of the CREB reaction. However, when CREB is present at saturating concentrations and in excess over β-catenin, Axin can produce larger rate increases for β-catenin phosphorylation. This effect arises because CREB can act as a competitive inhibitor of the β-catenin reaction. When Axin is added to the system, the *K*_M_ for CREB will increase, resulting in more GSK3β available to react with β-catenin and increasing the rate of β-catenin phosphorylation (Figure 5A). To test this prediction, we measured pS45-β-catenin phosphorylation in a competitive reaction with excess, saturating pS133-CREB present. In this system, adding Axin produced a 20-fold increase in pS45-β-catenin phosphorylation, much larger than the 3-fold increase in the absence of pS133-CREB (Figure 5B). As predicted, this larger effect results entirely from a decrease in the reaction rate in the absence of Axin, where CREB competes for GSK3β and inhibits the β-catenin reaction. When Axin is added, the rates for β-catenin phosphorylation are similar in the presence and absence of CREB because Axin suppresses the interaction of GSK3β with CREB. Thus, while Axin alone has a modest effect on the β-catenin reaction rate (Figure 2), in the presence of competing substrates, which is likely to be relevant *in vivo*, Axin can produce much larger increases in β-catenin phosphorylation.

**Figure 5.**
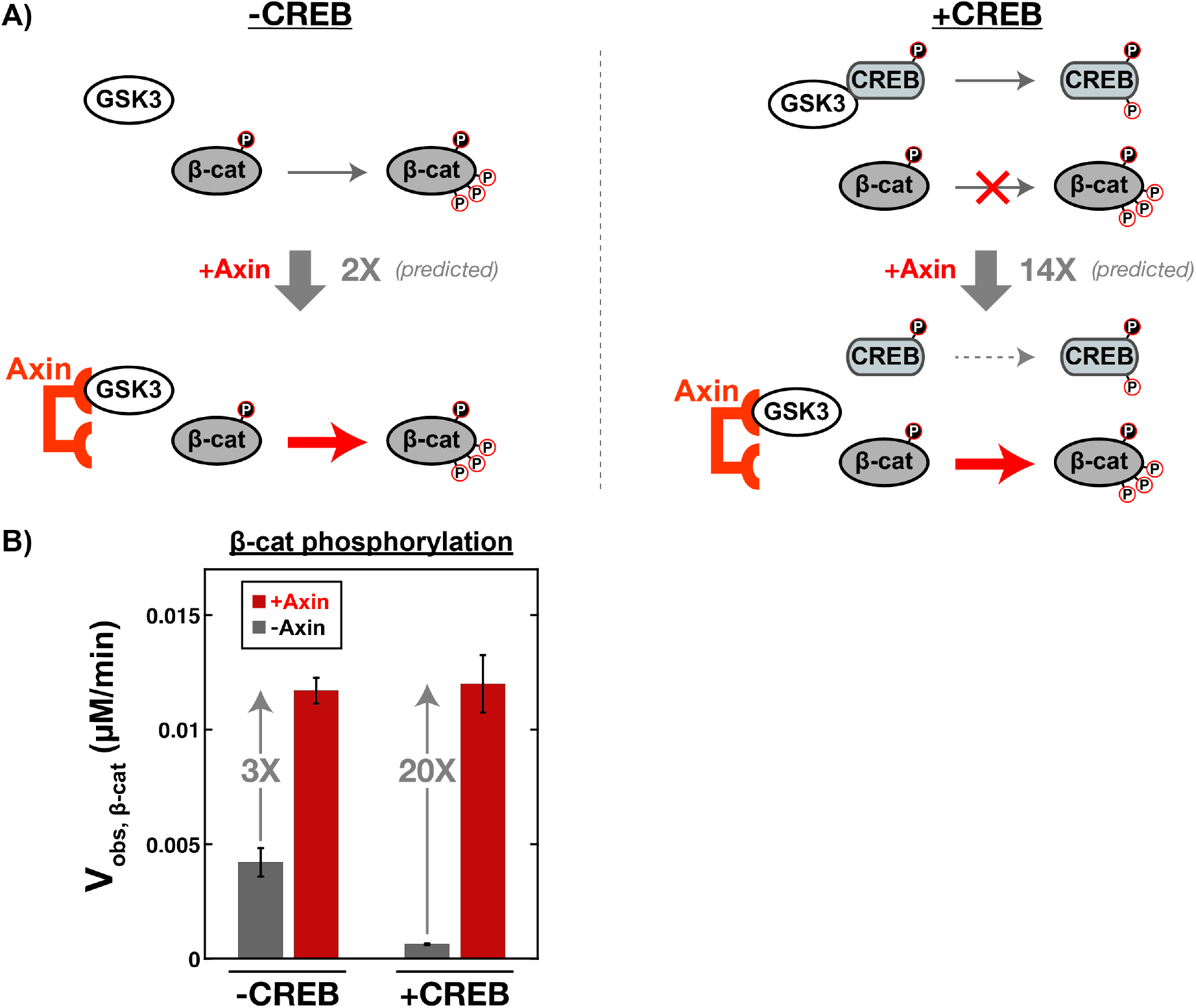
Axin accelerates the β-catenin reaction when competing substrates are present. (A) Prediction for the effect of Axin on pS45-β-catenin phosphorylation when the competing GSK3β substrate CREB is present. When pS133-CREB is present at saturating concentrations and pS45-β-catenin is present at subsaturating concentrations, pS133-CREB forms a complex with GSK3β and inhibits the reaction with pS45-β-catenin. Axin disrupts the interaction of GSK3β with CREB, preventing buildup of the GSK3β•CREB complex. Using a Michaelis-Menten model with CREB as a competitive inhibitor (Figure S6), we predict that Axin should produce a 14-fold increase on the β-catenin phosphorylation rate in the presence of CREB, much larger than the 2-fold increase in the absence of CREB. (B) β-catenin phosphorylation rates (*V*_obs_) measured at 20 nM GSK3β and 50 nM pS45-β-catenin (subsaturating) in the presence and absence of 500 nM miniAxin and/or 1 µM pS133-CREB (saturating). In the presence of CREB, Axin produces a 20-fold increase in the β-catenin phosphorylation rate. In the absence of CREB, Axin produces a 3-fold increase.

### An inactive GSK3β•β-catenin complex accumulates in the reaction with unprimed β-catenin

Our quantitative kinetic data described above suggest a new model for how the destruction complex specifically accelerates the β-catenin reaction, but a major question remains unanswered: if Axin has only ~2-fold effects on *k*_cat_ and *K*_M_ values for β-catenin phosphorylation, why did early biochemical studies in this system observe much larger rate increases? One notable difference is that the initial biochemical studies on Axin were performed with unprimed β-catenin as a substrate (18–20), prior to or concurrent with the discovery of phosphopriming by CK1 (21, 22). An early mechanistic model for Axin suggested that its function might be to bypass the need for phosphopriming (32). In this case, either phosphopriming of β-catenin or Axin-mediated tethering would promote the reaction of GSK3β with β-catenin, but there might not be an additional effect from Axin when β-catenin is phosphoprimed. When we measured reaction rates with unprimed β-catenin, we found that the unprimed reaction was ~10^2^-fold slower than the primed reaction, as expected. However, the steady state rate constants for unprimed β-catenin were not significantly affected by the presence of Axin (Figure S7, Table 1).

Additional experiments with unprimed β-catenin revealed an unexpected behavior that can explain previous reports of large Axin-mediated rate enhancements. In most circumstances, enzymatic reaction rates should increase linearly with increasing enzyme concentration, and this behavior occurs in the reactions with pS45-β-catenin and pS133-CREB (Figure 2C & 4C). However, when we varied the GSK3β concentration in reactions with unprimed β-catenin, we observed that the rate does not increase linearly with GSK3β concentration. Instead, in the absence of Axin the observed rates level off sharply above ~100 nM GSK3β. In contrast, in the presence of Axin the observed rates increase linearly with GSK3β concentration (Figure S8). This concentration-dependent inactivation of GSK3β could be due to the formation of an inactive dimer or oligomer, and Axin binding to GSK3β could prevent the formation of this inactive state. Thus, if reaction rates are measured at relatively high levels of GSK3β and with unprimed β-catenin, which was the case in the early biochemical studies (18–20), Axin produces a large increase in the observed rates.

Based on the kinetic data, the inactive state is likely a reversible, oligomeric GSK3β•β-catenin complex (Figures S8-S10). If the inactive state were a dimer or oligomer of GSK3β alone, we would have expected to see a similar inactivation effect in all GSK3β reactions, but this effect is not observed in reactions with pS45-β-catenin (Figure 2C) or with pS133-CREB (Figure 4C). Thus, the inactive state includes both GSK3β and unprimed β-catenin. Kinetic modeling suggests that the inactive state is likely a higher-order oligomer. The observed rates level off too sharply with increasing GSK3β concentration to fit to a dimer model (Figure S8). While the physiological relevance of an oligomeric, inactive GSK3β•β-catenin complex is uncertain (see discussion with Figure S10), it is important to recognize that the observation that Axin accelerates the β-catenin reaction *in vitro* was the initial foundation to explain the function of the Wnt pathway destruction complex *in vivo* (8–13). Our data suggest that the mechanistic origin of this original observation is only applicable to the specific condition of high GSK3β concentrations reacting with unprimed β-catenin, which is likely not relevant to the preferred physiological reaction of GSK3β with CK1-phosphoprimed pS45-β-catenin.

## Discussion

To understand how Wnt signals regulate Axin to modulate GSK3β activity, we biochemically reconstituted GSK3β-mediated reactions *in vitro* and attempted to reproduce the long-standing result that Axin substantially accelerates the reaction of GSK3β with β-catenin (18–20). Although prior reports suggested large effects as high as 10^4^-fold, we found modest 2- to 3-fold effects from Axin on the rate constants for the phosphorylation reaction (Table 1). Two key features of our work can explain this discrepancy. First, we measured *in vitro* reaction rates with the phosphoprimed form of β-catenin, while most prior work used unprimed β-catenin.^*^ A large Axin-dependent rate enhancement can be observed with unprimed β-catenin, but only in the specific condition of high GSK3β concentration, where Axin prevents the formation of an oligomeric, inactive GSK3β•β-catenin complex (Figure S8). Second, we measured well-defined rate constants in the presence and absence of Axin, while previous reports made indirect comparisons between observed rates, leading to the estimate of a 10^4^-fold effect from Axin (20).

The small effect of Axin on GSK3β activity towards β-catenin is the result of two opposing effects. First, Axin binding to GSK3β nonspecifically disrupts substrate binding, as seen in the reactions with CREB in the presence of Axin and with β-catenin in the presence of Axin∆BCD (Figures 3 & 4). Second, the β-catenin binding site allows Axin to rescue substrate binding specifically for β-catenin. These opposing effects produce the modest 2-fold decrease in *K*_M_ when comparing the β-catenin reactions of Axin•GSK3β to GSK3β, but a larger >7-fold decrease when comparing Axin•GSK3β to Axin∆BCD•GSK3β. Thus, it is reasonable to view Axin as a tethering scaffold that specifically promotes binding of GSK3β to β-catenin, even though the net effect of Axin on *K*_M_ is only 2-fold.

How do we reconcile the finding that Axin has small effects on reaction rates *in vitro* with the vast literature that supports the importance of Axin in Wnt signaling (11–13)? We know that Axin has substantial effects on Wnt signaling and vertebrate development (34–36), and Axin mutants are associated with cancer (37, 38). Moreover, the idea that the destruction complex acts to accelerate β-catenin phosphorylation is a cornerstone of functional models for Wnt signaling (16, 39). One important point to consider is that we do not have a clear framework to evaluate how large of an *in vitro* effect is necessary to account for phenotypic effects *in vivo*, and a 2-fold increase in reaction rates might actually be physiologically significant. In cell culture models, treatment with high levels of Wnt ligand can produce ~5-10-fold increases in total β-catenin levels (15, 40), with a corresponding ~5-fold decrease in GSK3β-β-catenin phosphorylation rate (15). Smaller changes can have significant effects *in vivo*, and transcriptional responses can be detected following ~2-fold changes in β-catenin levels (41). However, the observation that *in vivo* changes in β-catenin levels and phosphorylation rate can significantly exceed 2-fold suggests that we should be able to identify mechanisms that produce correspondingly larger effects *in vitro*.

While Axin alone does not appear to have a sufficiently large effect on β-catenin phosphorylation *in vitro* to account for *in vivo* observations, additional proteins and components that are present *in vivo* could lead to larger effects. We demonstrate one such possible contribution here: Axin produces >10-fold increases in reaction rates when another GSK3β substrate is present at saturating concentrations. The β-catenin reaction with GSK3β is inhibited by the presence of the competing substrate CREB, and Axin relieves this inhibition to produce a 20-fold increase in the β-catenin phosphorylation rate (Figure 5). This effect arises because Axin nonspecifically disrupts GSK3β binding to its substrates while simultaneously binding to β-catenin to maintain this specific interaction with GSK3β. In a cellular environment, where many competing substrates of GSK3β are present, free GSK3β could be saturated with non-Wnt pathway substrates and unable to react with β-catenin. An Axin-bound pool of GSK3β could preferentially phosphorylate β-catenin at rates much faster than the free, Axin-independent pool, leading to efficient β-catenin degradation. Wnt signals could prevent β-catenin phosphorylation by disrupting binding interactions to Axin or by perturbing the conformation of Axin to prevent effective tethering of β-catenin to GSK3β. Regardless of the precise mechanism, this model is consistent with the observation that GSK3β-dependent signaling pathways are insulated from each other (2–4). Measured cellular Axin concentrations are 10 to 10^3^-fold lower than GSK3β (39, 42–45), which means that only a small fraction of the total GSK3β can be bound to Axin. Thus, even if a Wnt signal disrupts the Axin•GSK3β complex and relieves the non-specific inhibition of GSK3β, the amount of GSK3β released into the Axin-independent pool would be small relative to the total amount of GSK3β, and would likely not significantly perturb phosphorylation rates towards other non-Wnt GSK3β substrates.

Additional components from within the Wnt pathway could also play an important role in promoting β-catenin phosphorylation in the destruction complex. APC is a particularly intriguing candidate (Figure 1B), as it binds Axin and has multiple binding sites for β-catenin in the low nM affinity range (46, 47). By binding tightly to β-catenin, the Axin•APC complex could be more effective than Axin alone at tethering β-catenin to GSK3β (48–50). Post-translational modifications of Axin and the formation of higher-order assemblies could also make significant contributions (16, 51). Finally, it is possible that the destruction complex also promotes the CK1α-mediated phosphopriming step (21, 22), which would increase the GSK3β-mediated β-catenin phosphorylation rate. There are conflicting reports on whether Axin accelerates the CK1α-catalyzed reaction *in vitro* (21, 33, 52), and there is evidence that Wnt signals affect both the CK1α and GSK3β-mediated reactions *in vivo* (15). An important feature of our work is that we have established a clear and quantitative framework to evaluate the functional effects of Axin on β-catenin phosphorylation, and we can now introduce additional components to evaluate their functional effects.

In addition to providing new insights into Wnt pathway signaling, our results have broad implications for understanding scaffold protein function. Axin was among the earliest proteins identified as a signaling scaffold (18), and was initially discussed as a prototypical model for tethering a kinase and substrate together to accelerate a phosphorylation reaction (53). There is now an emerging consensus that scaffold proteins can have many functions beyond simply tethering a kinase to its substrate, including allosterically modulating the activity of their target proteins (5). Here we show that Axin does have a tethering function, but the opposing effect from nonspecifically perturbing GSK3β binding to its substrates results in a modest net effect when the activity of Axin is studied in isolation. The functional advantage from Axin-mediated tethering only emerges in a more complex system with multiple competing substrates, which may more accurately reflect the *in vivo* environment. These insights arose from reconstitution and quantitative kinetic characterization of a minimal biochemical system *in vitro*, which highlighted an apparent discrepancy between *in vitro* and *in vivo* behavior and enabled us to identify a possible solution. Future biochemical studies that introduce additional Wnt pathway and GSK3β-interacting proteins will likely provide further insights and rigorous tests to expand our understanding of complex, interconnected cell signaling networks.

## Materials and Methods

### Protein Expression Constructs

The human Wnt pathway proteins GSK3β, β-catenin, and Axin (hAxin1 isoform 2, Uniprot O15169-2) were cloned into *E. coli* expression vectors containing an N-terminal maltose binding protein (MBP) and a C-terminal His6 tag. Human coding sequences were obtained as follows: GSK3β (addgene #14753)(54), β-catenin (addgene #17198, a gift from Randall Moon), and Axin (derived from hAxin1-rLuc, a gift from Randall Moon). CK1α was cloned with an N-terminal GST tag and a C-terminal His6 tag; the human CK1α sequence was obtained from addgene #92014 (55). The catalytic subunit of mouse PKA was expressed from pET15b with an N-terminal His-tag (addgene #14921)(56). The CREB_127-135_ peptide ILSRRPSYR was cloned by oligo annealing into a vector with an N-terminal MBP and C-terminal His6 tag. The coexpression plasmid for β-catenin and CK1α was constructed by inserting the GST-CK1α expression cassette (without the His6 tag) into the MBP-β-catenin-His plasmid. Axin truncation constructs Axin_384-518_ (miniAxin) and Axin_∆465-518_ (Axin∆BCD) were cloned as described for full length Axin above. MiniAxin contains binding sites for both GSK3β and β-catenin. The N-terminal boundary was defined based on the crystal structure of GSK3β with an Axin peptide (20) and the C-terminal boundary was defined based on the Pfam annotation of the BCD (PF08833) (57). The N-terminal boundary of the BCD (for Axin∆BCD) was also defined from the Pfam annotation. A complete list of protein expression constructs is provided in Table S2.

### Protein Expression and Functional Assays

Complete descriptions of purification methods, quantitative kinetic assays, and binding assays are included in the Supplementary Methods.

## Acknowledgements

We thank Dustin Maly, Dan Herschlag, John Scott, Jonathan Cooper, David Kimelman, Wendell Lim, Michael Gelb, Steven Wiley, Geeta Narlikar, Wenqing Xu, Renée van Amerongen, Edmond Fischer, and members of the Zalatan group for comments and discussion. We thank Sujata Chakraborty for purifying YopH. This work was supported by a Career Award at the Scientific Interface from the Burroughs Wellcome Fund (J.G.Z.) and NIH R35 GM124773 (J.G.Z.).

## Author contributions

M.G., E.F., E.B.S., and J.G.Z. designed research; M.G., E.F., and E.B.S. performed research; and M.G., E.F., E.B.S., and J.G.Z. wrote the paper.

## Footnotes

* One study included both CK1 and GSK3β in a reaction with β-catenin in the presence and absence of Axin (33). The effect of Axin on the observed rates reported in that work is consistent with the rate constants we observe here.

