## Supplementary Information for "The Wnt pathway scaffold protein Axin promotes signaling specificity by suppressing competing kinase reactions"

**This PDF file includes:**

Supplementary methods
Figures S1 to S10
Tables S1 to S2
SI References

### Supplementary Methods

*Protein Expression and Purification*

All Wnt pathway, CREB, and PKA proteins were expressed in Rosetta (DE3) pLysS *E. coli* cells by inducing with 0.5 mM IPTG overnight at 18 ºC. Constructs with N-terminal MBP and C-terminal His6 tags (GSK3β, β-catenin, Axin, and $CREB_{127-135}$) were affinity purified with HisPur Ni-NTA resin (Thermo Scientific) and amylose resin (NEB). CK1α was purified with Ni-NTA resin and glutathione agarose resin (Thermo Scientific). The PKA catalytic subunit was purified on Ni-NTA resin. Purified proteins were dialyzed into 20 mM Tris-HCl pH 8.0, 150 mM NaCl, 10% glycerol, and 2 mM DTT at 4 °C, aliquoted and stored at -80 °C. If necessary, proteins were concentrated using 10000 or 30000 MWCO Amicon Ultra-15 Centrifugal Filter devices at 4 °C, 2000×g. Protein concentrations were determined using a Bradford assay (Thermo Scientific). YopH was expressed in BL21 (DE3) *E. coli* cells and purified as previously described (1).

For quantitative binding assays using bio-layer interferometry, GSK3β was further purified by size exclusion chromatography using a Superdex 200 Increase 10/300 GL column (GE Healthcare) and used immediately.

*Quantitative Western Blotting*

Protein samples were run on 4-15% miniProtean TGX gels (Bio-Rad) and transferred to 0.2 μm nitrocellulose membranes (Bio-rad). Membranes were blocked with 50:50 Li-Cor blocking buffer:TBST, incubated with antibodies as described below using either manual washes or a Precision Biosystems BlotCycler, visualized using the Li-Cor Odyssey Imaging System, and analyzed using Image Studio Lite 5.2.5 (Li-Cor).

*Determination of the phosphorylation state of GSK3β Tyr216*

200 nM purified GSK3β was incubated with 12 μM tyrosine phosphatase (YopH) for 30 minutes in PMP buffer (New England Biolabs) [50 mM HEPES pH 7.5, 100 mM NaCl, 2 mM DTT, 0.01% Brij 35] at 25 ºC. The level of Tyr216 phosphorylation was assessed by western blotting using a primary Anti-GSK-3β (pY216) antibody (BD Biosciences #612312) and a secondary IRDye 800CW Donkey Anti-Mouse IgG antibody (Li-Cor #926-32212). Total GSK3β (MBP-tagged) was detected by western blotting using a primary MBP Tag (8G1) antibody (Cell Signaling Technology #2396) and a secondary IRDye 800CW Donkey Anti-Mouse IgG antibody (Li-Cor #926-32212).

*Preparation of phospho-primed β-catenin (pS45-β-catenin)*

For quantitative kinetic and binding assays, pS45-β-catenin was obtained by co-expression with CK1α in *E. coli* and purified as described above for β-catenin.

pS45-β-catenin could also be obtained by *in vitro* phosphorylation of β-catenin with purified CK1α. The phosphorylation reaction was performed in a 5 mL volume with 5 μM β-catenin, 1 μM CK1α, and 500 μM ATP in 40 mM HEPES pH 7.4, 50 mM NaCl, 10 mM $MgCl_2$, 0.05% IGEPAL. The reaction was incubated at 25 ºC for 1.5 hours.

For both co-expressed and *in vitro* phosphorylated pS45-β-catenin, the extent of pS45 phosphorylation was evaluated by western blotting using a primary anti-phospho-β-Catenin (Ser45) antibody (Cell Signaling Technology #9564) and a secondary IRDye 800CW Goat Anti-Rabbit IgG antibody (Li-Cor #926-32211).

*Preparation of phospho-primed CREB (pS133-CREB$_{127-135}$)*

Preparative-scale *in vitro* phosphorylation of CREB$_{127-135}$ at Ser133 was performed in a 10 mL reaction with 13 μM CREB$_{127-135}$, 6.5 μM PKA, and 100 μM ATP in 40 mM HEPES pH 7.4, 50 mM NaCl, 10 mM $MgCl_2$, 0.05% IGEPAL. The reaction was incubated at 25 ºC for 20 minutes. pS133-CREB$_{127-135}$ was purified with amylose resin, concentrated in storage buffer [20 mM Tris-HCl pH 8.0, 150 mM NaCl, 10% glycerol, and 2 mM DTT], aliquoted and stored at -80 °C.

To determine the time necessary for preparative-scale CREB phosphorylation, an analytical-scale phosphorylation reaction was performed with γ-$^{32}$P-ATP. A 30 μL reaction was conducted with 10 μM CREB$_{127-135}$ and 2 μM PKA in 40 mM HEPES pH 7.4, 50 mM NaCl, 10 mM $MgCl_2$, 0.05% IGEPAL at 25 ºC. The reaction was initiated by adding ATP to a final concentration of 100 μM unlabeled ATP and 0.24 μCi γ-$^{32}$P-ATP. Timepoints were collected at 5, 10, 20, 40, and 60 minutes. 5 μL aliquots were quenched by spotting on 0.2 μm nitrocellulose membranes (Bio-rad) and placing the membranes in 0.5% v/v phosphoric acid. Membranes were washed 4X 5 min in 0.5% v/v phosphoric acid, allowed to air dry, exposed to a phosphorimager cassette, and imaged on a GE Typhoon FLA 9000. Images were analyzed using ImageQuant 5.1 (GE Healthcare). The data indicated that the reaction approached completion within the first 5-10 minutes (Figure S5). We performed preparative scale CREB phosphorylation reactions using a similar concentration of CREB and >3X more PKA, which should go to completion in <20 minutes.

*Quantitative Kinetic Assays*

*In vitro* kinetic assays were conducted in kinase assay buffer [40 mM HEPES pH 7.4, 50 mM NaCl, 10 mM $MgCl_2$, and 0.05% IGEPAL] at 25 ºC in 60 μL total volume. Reactions were initiated by adding ATP to a final concentration of 100 μM (this concentration of ATP is saturating for GSK3β – see Figure S3). Reaction timepoints for initial rate kinetics were obtained at 10, 30, 60, and 90 seconds (pS45-β-catenin reactions); 2, 5, 10, and 20 minutes (unprimed β-catenin reactions); 0.5, 1, 2, and 3 minutes (CREB reactions at [GSK3β] > 20 nM); and 2, 5, 10, and 15 minutes (CREB reactions at [GSK3β] = 20 nM). 10 μL aliquots were quenched by boiling in 5X SDS loading buffer. Samples were analyzed by SDS-PAGE and quantitative western blotting as described above. For reactions with [β-catenin] ≥ 500 nM, gel samples were diluted 2-fold (unprimed β-catenin reactions) or 5-fold (pS45-β-catenin reactions) to prevent a gel smearing artifact.

GSK3β-phosphorylated β-catenin was detected using a primary anti-Phospho-β-Catenin (Ser33/37/Thr41) Antibody (Cell Signaling Technology #9561). GSK3β-phosphorylated CREB was detected using a primary anti-Phospho-CREB (Ser129) Antibody (Thermo Scientific PA5-36843). For both products, the secondary antibody was IRDye 800CW Goat Anti-Rabbit IgG antibody (Li-Cor #926-32211).

Concentrations of phosphorylated β-catenin product were determined by comparing western blot signal intensities to an endpoint standard containing 50 nM β-catenin phosphorylated to completion with GSK3β. The standard was prepared in a reaction with 50 nM unprimed β-catenin, 100 nM GSK3β, 100 nM miniAxin, and 100 μM ATP in kinase assay buffer at 25 ºC for 17 hrs, or in a reaction with 50 nM pS45-β-catenin, 100 nM GSK3β, 100 nM miniAxin, and 100 μM ATP in kinase assay buffer at 25 ºC for 25 min. The signal intensities for endpoints prepared from unprimed and phosphoprimed β-catenin were indistinguishable.

Concentrations of phosphorylated CREB product were determined similarly with an endpoint standard containing 50 nM CREB phosphorylated to completion with GSK3β. The standard was prepared in a reaction with 50 nM pS133-CREB, 500 nM GSK, 100 μM ATP in kinase assay buffer at 25 ºC for 2.5 hrs.

Kinetic parameters were determined by fitting plots of initial rates ($V_{obs}$) vs. [substrate] to the Michaelis-Menten equation ($V_{obs} = k_{cat}[E]_0[S]/(K_M+[S])$). For the reaction of CREB in the presence of Axin, which did not detectably saturate, the value of $k_{cat}/K_M$ was obtained from the slope of a linear fit to a plot of $V_{obs}$ vs. [substrate]. For the inactive GSK3β oligomer model (Figure S8), the data were fit to the equation: $[GSK3\beta]_{total} = (V_{obs}/k_{cat}) + (N/K_{oligomer})\times(V_{obs}/k_{cat})^N$.

*Quantitative Binding Assays*

Binding affinities ($K_D$) and the corresponding association and dissociation rate constants ($k_a$ and $k_d$) were determined with bio-layer interferometry using an Octet Red96e system (ForteBio) and Streptavidin biosensor tips. Proteins were biotinylated using an EZ-Link Micro NHS-PEG4-Biotinylation kit (Thermo Scientific) with a molar coupling ratio of 1:1 and purified using 7K MWCO Zeba Spin Desalting Columns (Thermo Scientific). Binding assays were performed in 10 mM $Na_2HPO_4/NaH_2PO_4$ pH 7.4, 137 mM NaCl, 2.6 mM KCl, 0.1% BSA, and 0.02% Tween-20 at 22 °C with Greiner Bio-One 96-Well Non-treated Polypropylene Microplates (Fisher Scientific) at a shake speed of 1000 rpm. Well volumes were 200 μL. Background buffer effects were corrected using a buffer reference sample (a tip loaded with biotinylated protein and dipped into buffer instead of analyte). The buffer reference sample was subtracted from the binding assay data before analysis. Data were analyzed using Data Analysis HT 11.0 (ForteBio) to obtain values of $K_D$, $k_a$ and $k_d$.

For the binding interaction between GSK3β and miniAxin, the miniAxin protein was biotinylated. Biosensor tips were hydrated for 10 minutes in assay buffer, equilibrated for 1 minute, and then loaded with 10 nM biotinylated miniAxin for 5 minutes. After a 1 minute equilibration in assay buffer, tips were immersed in varying concentrations of GSK3β (500 nM, 250 nM, 125 nM, 62.5 nM, 31.3 nM, 15.6 nM, and 7.81 nM) for 5 minutes to monitor association kinetics. Tips were then immersed in assay buffer for 10 minutes to measure dissociation kinetics.

For the binding interaction between pS45-β-catenin and miniAxin, pS45-β-catenin was biotinylated. Biosensor tips were hydrated for 10 minutes in assay buffer, equilibrated for 1 minute, and then loaded with 30 nM nM biotinylated pS45-β-catenin for 5 minutes. Tips were sequentially washed 3X for 1 minute in assay buffer to minimize a baseline drift effect that occurred if these wash steps were omitted. Tips were immersed in varying concentrations of miniAxin (10 μM, 2.5 μM, 625 nM, 156.25 nM) for 100 seconds to monitor association kinetics. Tips were then immersed in assay buffer for 60 seconds to measure dissociation kinetics.

### Figure S1

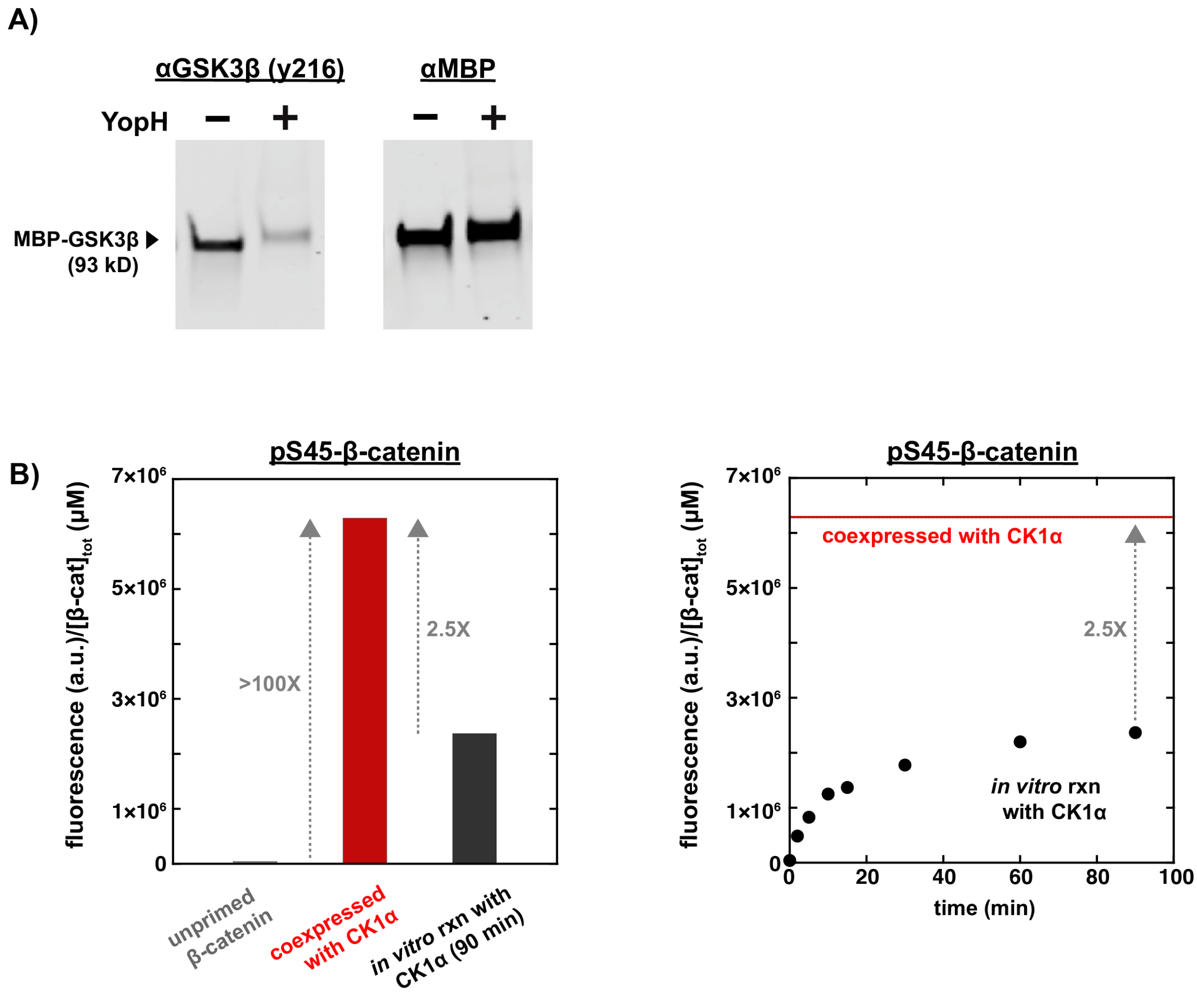


**Figure S1.** (A) GSK3β purified from *E. coli* is phosphorylated on the activation loop Tyr216. Purified GSK3β was incubated with the tyrosine phosphatase YopH and analyzed by western blot using an antibody specific for pY216 GSK3β. Purified GSK3β produced a detectable signal at the expected molecular weight (94 kD), and the signal decreases substantially upon YopH treatment. The loading control indicates no change in total GSK3β, using an anti-MBP antibody that detects the MBP tag on purified GSK3β. (B) β-catenin co-expressed with CK1α in *E. coli* is phosphorylated on Ser45. The extent of S45 phosphorylation was detected by western blot using an antibody specific for pS45-β-catenin. Phosphoprimed β-catenin can also be obtained by incubating unprimed β-catenin with purified CK1α. In this reaction, we observe partial phosphorylation of Ser45, with a total pS45 signal (normalized to total protein concentration) reaching only ~40% of that obtained when β-catenin is co-expressed with CK1α.

### Figure S2

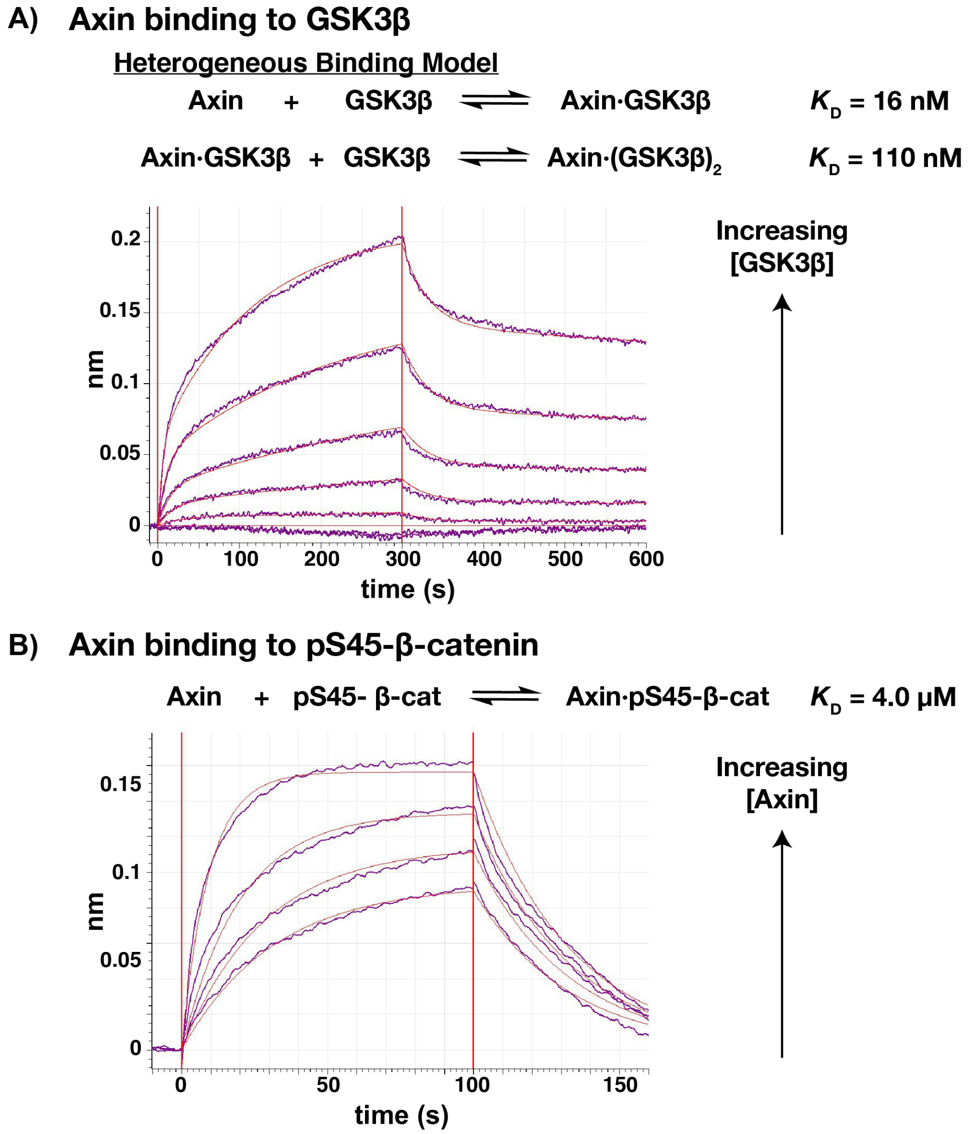


**Figure S2.** Axin binding to GSK3β and β-catenin. (A) Bio-layer interferometry traces showing association and dissociation of GSK3β to immobilized biotinylated miniAxin. Similar data were obtained with full length Axin and AxinΔBCD. The data were fit to a Heterogeneous Binding Model (Data Analysis HT 11.0, ForteBio) that accounts for GSK3β binding to Axin and GSK3β dimerization, which has been previously reported (2). This fit gives a $K_D$ of 16 nM for miniAxin binding to GSK3β. The fitted $K_D$ for GSK3β dimerization is 110 nM, which is somewhat tighter than the low μM value estimated from crosslinking experiments (2). A 1:1 binding model that neglects GSK3β dimerization gives a poor fit to the data. Values of fitted binding constants are reported in Table 1. (B) Bio-layer interferometry traces showing association and dissociation of miniAxin to immobilized biotinylated pS45-β-catenin. This binding event produced a negative shift in the interferometry signal (the data are displayed with an inverted y-axis), indicating a compression in the surface which could result from a conformational change in the bound complex. The data were fit using a 1:1 binding model. This fit gives a $K_D$ of 4.0 μM for Axin binding to pS45-β-catenin.

**Figure S3**

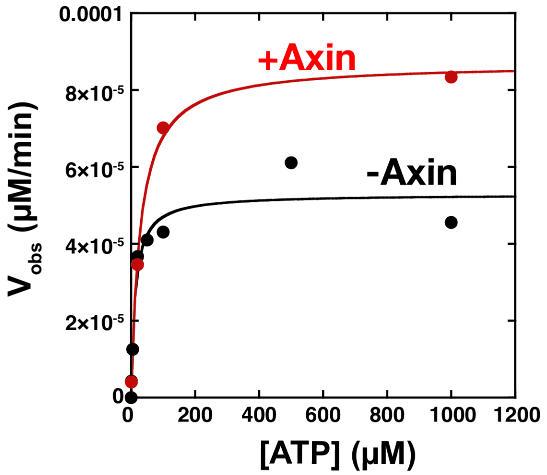


**Figure S3.** The concentration of ATP used for quantitative kinetic experiments (100 µM) is saturating for GSK3β in the presence and absence of Axin. Michaelis-Menten plot of $V_{obs}$ vs. [ATP] at 20 nM GSK3β and 500 nM unprimed β-catenin in the presence and absence of 100 nM miniAxin. Fits to the Michaelis-Menten equation give $K_{M,\,ATP}$ values of 12 ± 4 µM and 28 ± 3 µM in the presence and absence of Axin, respectively.

**Figure S4**

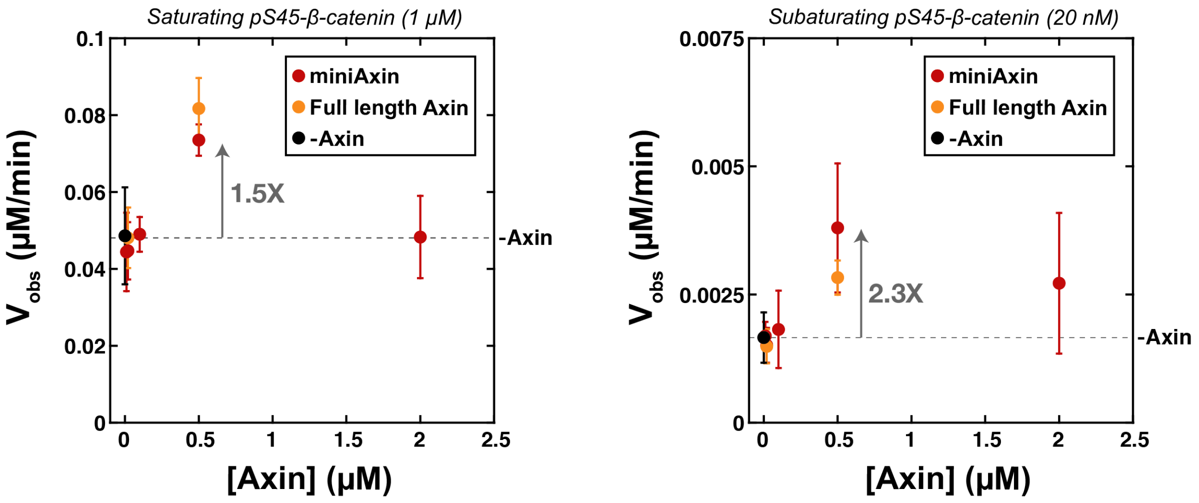


**Figure S4.** Varying the concentration of Axin for the reaction of GSK3β with pS45-β-catenin does not produce larger rate effects than observed with 500 nM Axin (Figure 2). Plots of $V_{obs}$ vs. [Axin] with either miniAxin or full length Axin, 20 nM GSK3β, and 1 μM or 20 nM pS45-β-catenin. It is important to test both saturating concentrations (i.e. $k_{cat}$ conditions, 1 μM pS45-β-catenin) and subsaturating concentrations (i.e. $k_{cat}/K_M$ conditions, 20 nM pS45-β-catenin) because if Axin has a large effect on $K_M$ it would not be detectable in saturating conditions. For full length Axin, only a subset of Axin concentrations were measured (20 and 500 nM). Error bars are mean ± SD for at least 3 measurements.

### Figure S5

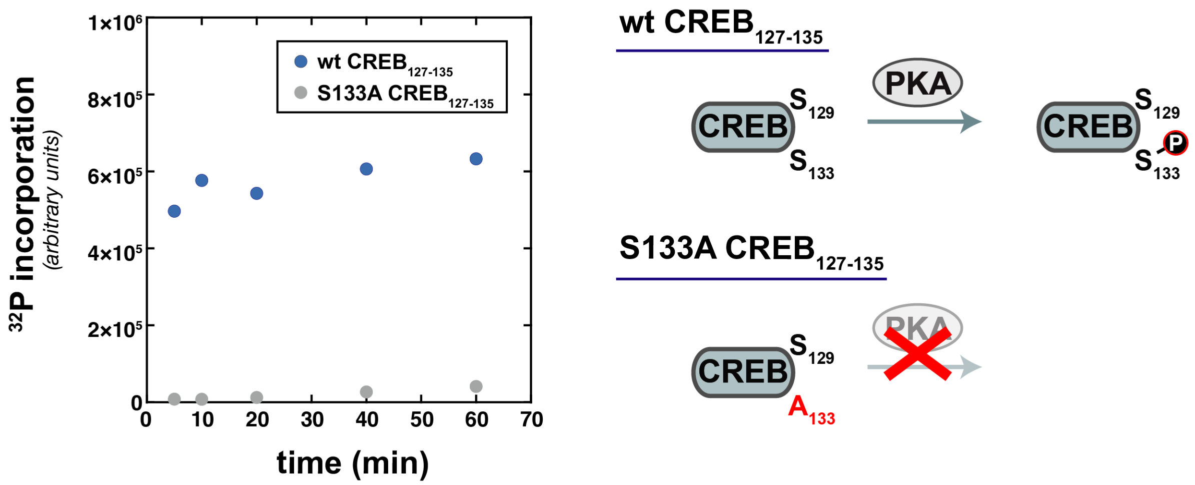


**Figure S5.** PKA phosphorylates $CREB_{127\text{-}135}$ on $Ser_{133}$. Reactions were performed with 10 µM $CREB_{127\text{-}135}$, 2 µM PKA, 100 µM ATP, and trace γ-$^{32}$P-ATP as described in the Methods section. $^{32}$P incorporation into $CREB_{127\text{-}135}$ rapidly approaches completion within the first 5-10 minutes of the reaction. No significant $^{32}$P incorporation is detectable in S133A $CREB_{127\text{-}135}$, which indicates that wt $CREB_{127\text{-}135}$ is phosphorylated on $Ser_{133}$.

### Figure S6

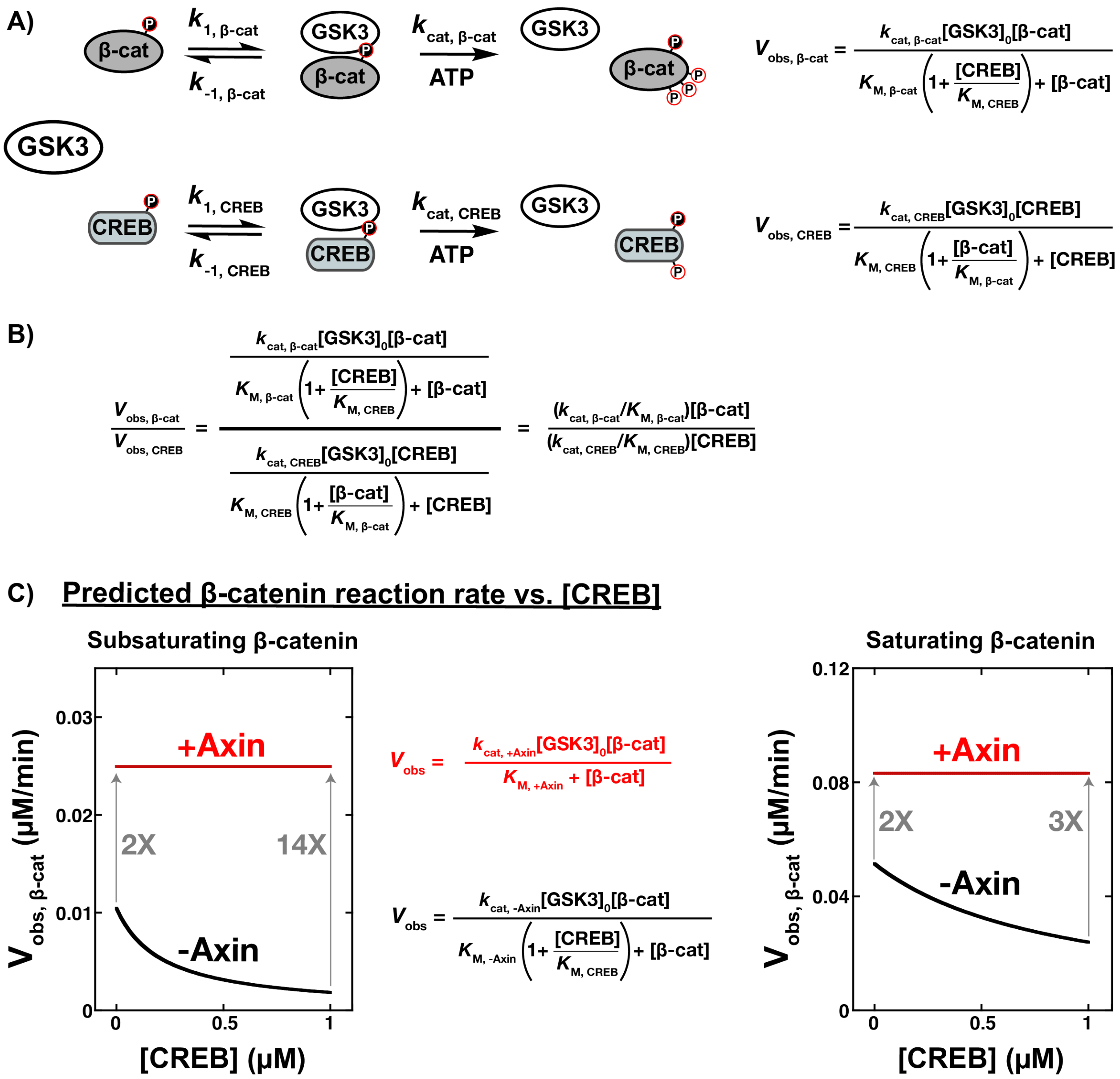


**Figure S6.** Kinetic model for a competitive reaction between β-catenin and CREB. (A) Kinetic scheme for a competitive reaction. Each substrate effectively acts as a competitive inhibitor for the other substrate. (B) The rate equations reduce to the expected specificity equation, where the ratio of the rates is determined by the ratios of $k_{cat}/K_M$ and the substrate concentrations (3). (C) Predicted rates for β-catenin phosphorylation as a function of CREB concentration at 20 nM GSK3β and 50 nM (subsaturating) or 1 µM (saturating) pS45-β-catenin. Increasing concentrations of CREB inhibit the β-catenin reaction in the absence of Axin. The expected values of $V_{obs}$ were calculated using the equations from (A) and the kinetic parameters from Table 1. For the +Axin reaction, where $K_{M,\ CREB} \geq 1$ µM, we assumed that the $[CREB]/K_{M,\ CREB}$ term is negligible over the [CREB] concentration range shown. When β-catenin is present at saturating concentrations, CREB is less effective at competing for GSK3β, leading to a smaller effect from Axin at high [CREB].

**Figure S7**

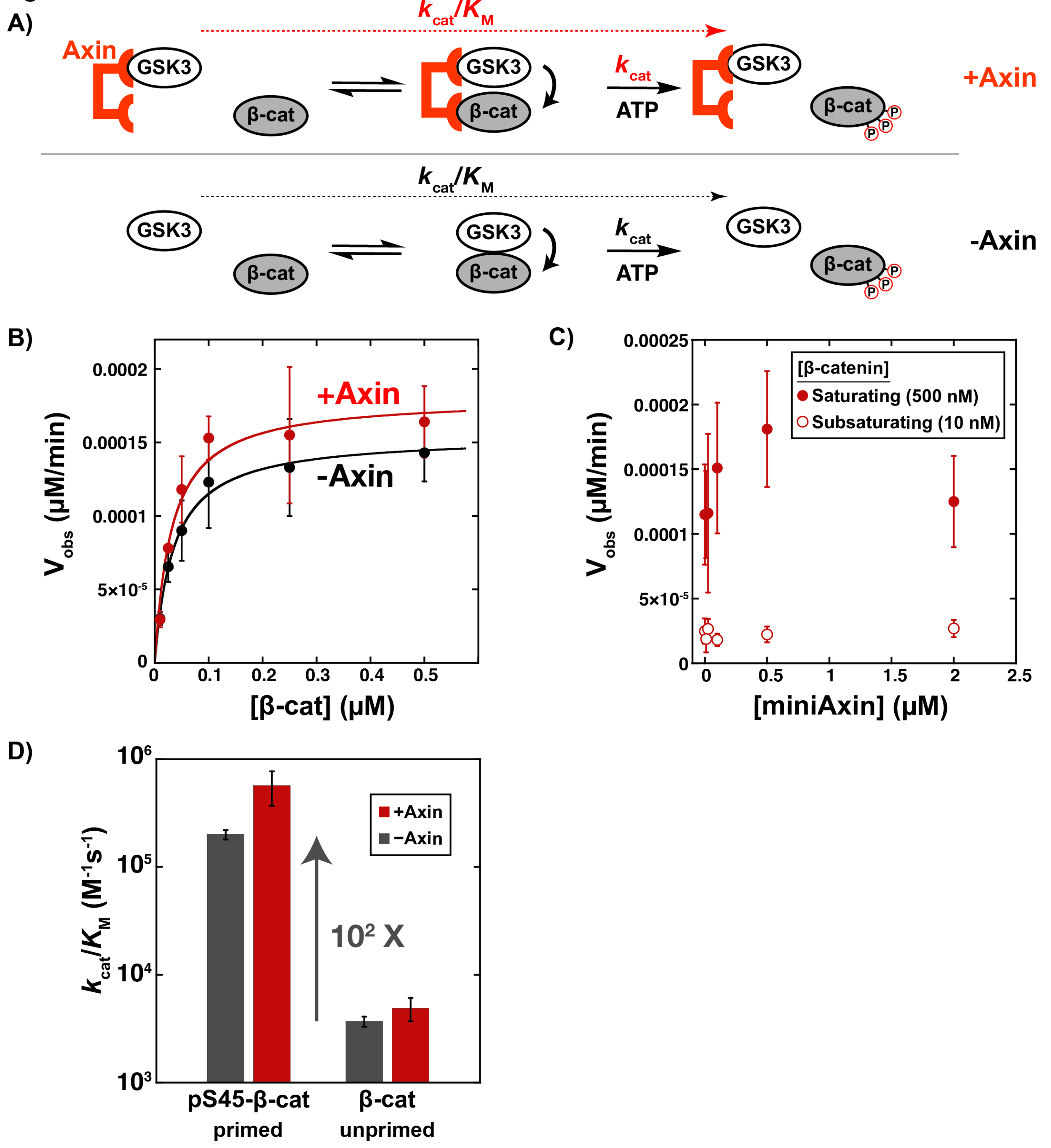


**Figure S7.** Axin has no significant effect on phosphorylation of unprimed β-catenin. (A) Minimal kinetic scheme for the reaction of GSK3β with β-catenin in the presence and absence of Axin, following Figure 2A. (B) Michaelis-Menten plot of $V_{obs}$ vs. [β-catenin] at 20 nM GSK3β in the presence and absence of 500 nM miniAxin. (C) Varying the concentration of Axin for the reaction of GSK3β with unprimed β-catenin does not produce significant rate effects. Plot of $V_{obs}$ vs. [miniAxin] at 20 nM GSK3β and 10 nM (subsaturating) or 500 nM (saturating) β-catenin. Error bars for (B) and (C) are mean ± SD for at least 3 measurements. (D) Comparison of $k_{cat}/K_M$ values (Table 1) for phosphoprimed and unprimed β-catenin.

### Figure S8

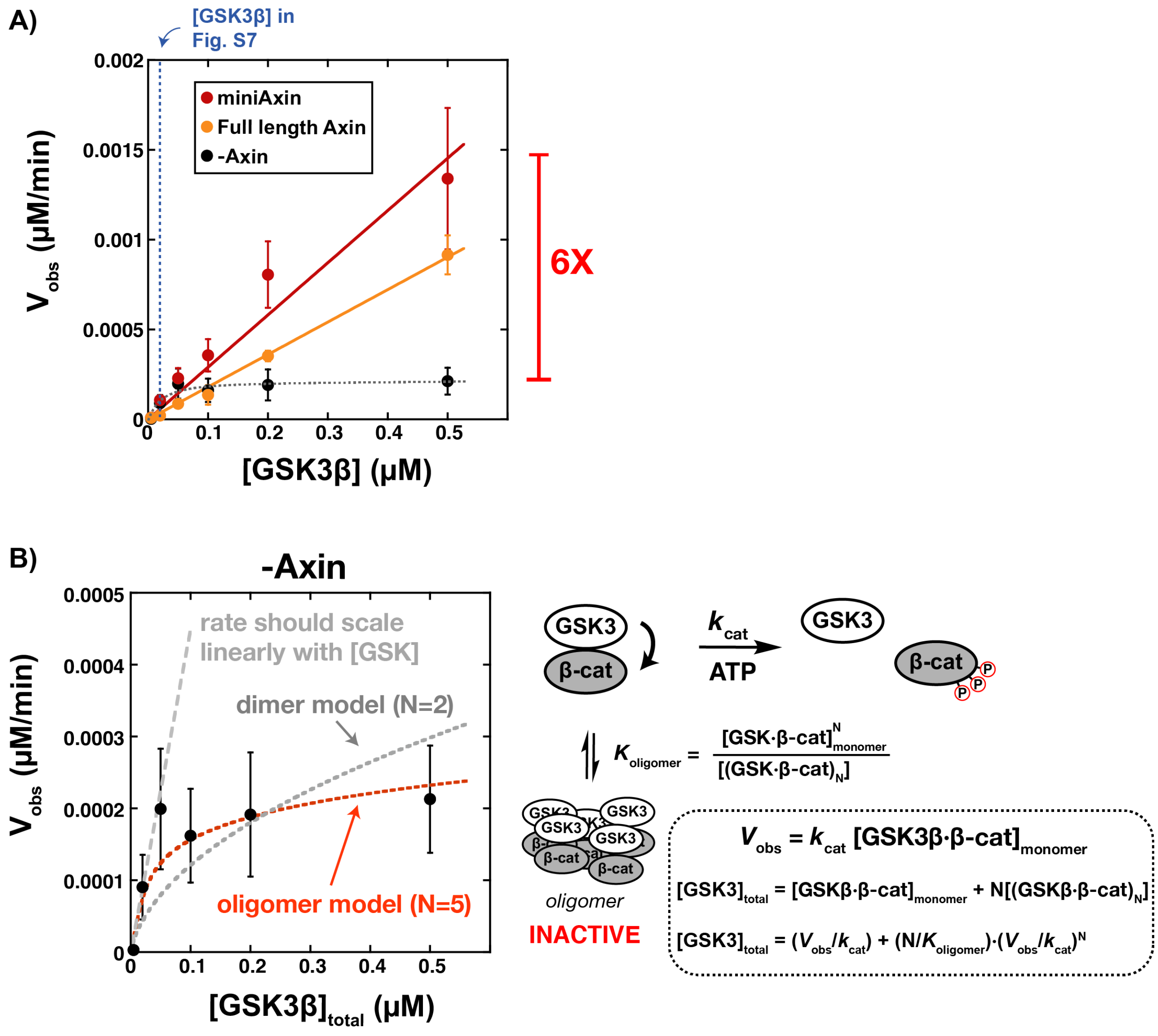


**Figure S8.** Axin rescues GSK3β from an inactive state. (A) Plot of $V_{obs}$ vs. [GSK3β] at 500 nM unprimed β-catenin in the presence and absence of 500 nM miniAxin or 500 nM full length Axin. (B) Expanded view of the plot of $V_{obs}$ vs. [GSK3β] in the absence of Axin, and a kinetic model with an inactive, oligomeric state. In the absence of the inactive state, $V_{obs}$ should increase linearly with [GSK3β]. Fitting the data to a model with an inactive GSK3β dimer (N=2) gives non-linear behavior but still shows substantial deviations from the data. Fitting the data to a model with a higher-order inactive GSK3β dimer (N=5) gives a reasonable fit. Larger values of N give indistinguishable fits to the data. The N=5 fit is also shown in panel (A). Error bars in both panels are mean ± SD for at least 3 measurements.

**Figure S9**

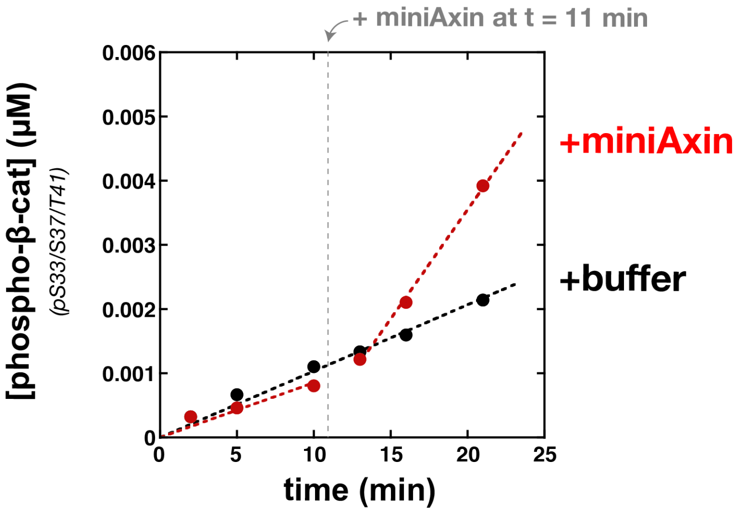


**Figure S9.** Formation of the inactive GSK3β•β-catenin complex is reversible. Product vs. time plot for a reaction with 500 nM GSK3β and 500 nM unprimed β-catenin. At t = 11 min, miniAxin was added to a final concentration of 500 nM and the reaction rate sharply increases. A control reaction in which an equal volume of buffer was added shows no change in rate. These data suggest that the inactive state is not an irreversible aggregate of GSK3β and unprimed β-catenin.

**Figure S10**

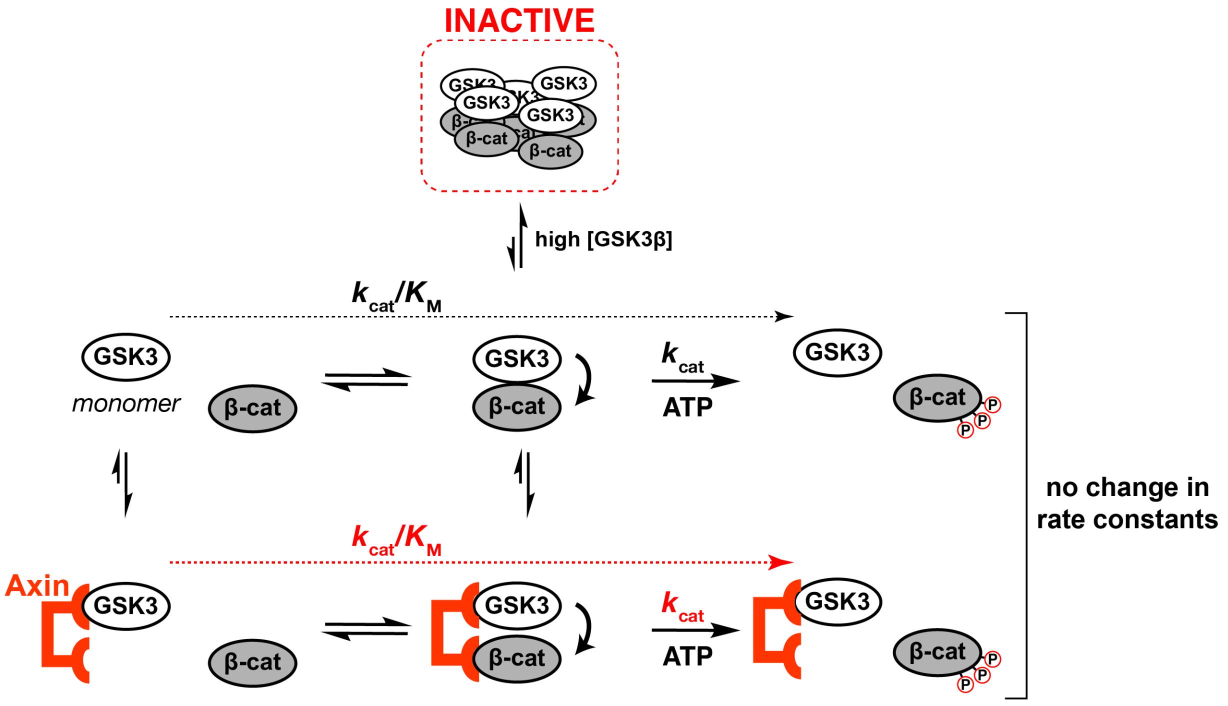


**Figure S10.** A revised kinetic model for the effect of Axin on the reaction of GSK3β with unprimed β-catenin. At low GSK3β concentrations, GSK3β is predominantly monomeric and active, and Axin has no significant effect on the steady state rate constants for the reaction (Figure S7). At high GSK3β concentrations, the GSK3β•β-catenin complex cooperatively assembles into an inactive, higher-order oligomer. Axin prevents the formation of this higher-order oligomer, which leads to an increase in observed rates for the reaction of GSK3β with β-catenin (Figure S8). The Axin-dependent rate enhancement at high GSK3β concentration arises because Axin increases active enzyme concentration, not because of a tethering effect between kinase and substrate.

The physiological relevance of an inactive, oligomeric GSK3β•β-catenin complex is uncertain. Cellular GSK3β concentrations estimated from mass spectrometry proteomics datasets (4-6) vary in the range of ~10-300 nM for human HeLa and U2OS cells (cell volumes from BioNumbers BNID 103725 & 108088 (7)). These values are similar to the ~100 nM concentration where we see the active/inactive transition *in vitro*, and additional experiments to determine if an oligomeric GSK3β•β-catenin complex exists *in vivo* could be justified.

### Supplementary Tables

**Table S1.** Binding constants for Axin interacting with GSK3β and β-catenin.[a]

| Immobilized Substrate | Binding Partner | $K_D$ | $k_a$ ($M^{-1}s^{-1}$) | $k_d$ ($s^{-1}$) |
|---|---|---|---|---|
| miniAxin | GSK3β | 16 nM | $1.7 \times 10^4$ | $2.7 \times 10^{-4}$ |
| Axin (full length) | GSK3β | 7.5 nM | $2.7 \times 10^4$ | $2.0 \times 10^{-4}$ |
| AxinΔBCD | GSK3β | 11 nM | $1.9 \times 10^4$ | $2.1 \times 10^{-4}$ |
| pS45-β-catenin | miniAxin | 4.0 μM | $6.4 \times 10^3$ | $7.3 \times 10^{-2}$ |

[a] Binding constants were determined using bio-layer interferometry as described in the Methods. See Figure S2. All fits had $\chi^2$ values < 1 and $R^2$ values > 0.98.

**Table S2.** Protein expression plasmids.

| Plasmid | Protein[a] | Expressed Protein | Vector | Source |
|---|---|---|---|---|
| pEF019 | β-catenin | MBP-β-catenin-His | pMBP-MG[b] | *This study* |
| pES001 | GSK3β | MBP-GSK3β-His | pMBP-MG | *This study* |
| pEF073 | Axin | MBP-Axin-His | pMBP-MG | *This study* |
| pMG023 | $Axin_{384-518}$ (miniAxin) | MBP-$Axin_{384-518}$-His | pMBP-MG | *This study* |
| pMG035 | $Axin_{\Delta 465-518}$ (AxinΔBCD) | MBP-$Axin_{\Delta 465-518}$-His | pMBP-MG | *This study* |
| pMG046 | CK1α | GST-CK1α-His | pETARA | *This study* |
| pMG051[c] | β-catenin/CK1α | MBP-β-catenin-His/GST-CK1α | pMBP-MG | *This study* |
| pEF086 | CREB (127-135) | MBP-$CREB_{127-135}$-His | pMBP-MG | *This study* |
| $H_6$-rC | PKA catalytic subunit | His-PKA-rC | pET15b | Addgene #14921 |
| YopH | YopH | YopH (untagged) | pCDFDuet-1 | (1) |

[a] All proteins are human sequences except PKA, which is the mouse sequence.
[b] pMBP-MG is a modified version of pMAL-p2X (New England Biolabs) with an N-terminal TEV-cleavable MBP tag and a C-terminal His6 tag. pETARA contains an N-terminal TEV-cleavable GST tag and a C-terminal His6 tag. pMBP-MG and pETARA were described previously (8).
[c] pMG051 was constructed by inserting the GST-CK1α expression cassette (without the His tag) from pMG046 into the pEF019 backbone.
